# Arc mediates intercellular tau transmission via extracellular vesicles

**DOI:** 10.1101/2024.10.22.619703

**Authors:** Mitali Tyagi, Radhika Chadha, Eric de Hoog, Kaelan R. Sullivan, Alicia C. Walker, Ava Northrop, Balazs Fabian, Monika Fuxreiter, Bradley T. Hyman, Jason D. Shepherd

**Affiliations:** Department of Neurobiology, University of Utah, Salt Lake City, USA; Department of Theoretical Biophysics, Max Planck Institute of Biophysics, Germany; Department of Biomedical Sciences University of Padova, Padova, Italy; Department of Neurology, Massachusetts Alzheimer’s Disease Research Center, Massachusetts General Hospital, Harvard Medical School, Boston, USA

## Abstract

Intracellular neurofibrillary tangles that consist of misfolded tau protein^1^ cause neurodegeneration in Alzheimer’s disease (AD) and frontotemporal dementia (FTD). Tau pathology spreads cell-to-cell^2^ but the exact mechanisms of tau release and intercellular transmission remain poorly defined. Tau is released from neurons as free protein or in extracellular vesicles (EVs)^3–5^ but the role of these different release mechanisms in intercellular tau transmission is unclear. Here, we show that the neuronal gene Arc is critical for packaging tau into EVs. Brain EVs purified from human tau (hTau) transgenic rTg4510 mice (rTg^WT^) contain high levels of hTau that are capable of seeding tau pathology. In contrast, EVs purified from rTg^WT^ crossed with Arc knock-out mice (rTg^Arc^ ^KO^) have significantly less hTau and cannot seed tau aggregation. Arc facilitates the release of hTau in EVs produced via the I-BAR protein IRSp53, but not free tau. Arc protein directly binds hTau to form a fuzzy complex that we identified in both mouse and human brain tissue. We find that pathological intracellular hTau accumulates in neurons in rTg^Arc^ ^KO^ mice, which correlates with accelerated neuron loss in the hippocampus. Finally, we find that intercellular tau transmission is significantly abrogated in Arc KO mice. We conclude that Arc-dependent release of tau in EVs plays a significant role in intracellular tau elimination and intercellular tau transmission.

## Introduction

Neurodegenerative diseases are characterized by protein aggregation in specific brain regions that spread across the brain as the disease progresses. A major histopathological hallmark of Alzheimer’s disease (AD) and tauopathies, such as frontotemporal dementia (FTD and chronic traumatic encephalopathy^6^, is intracellular neurofibrillary tangles that consist of misfolded tau protein^1^. Tau is a microtubule-associated protein that normally regulates microtubules to ensure proper cytoskeletal organization and cargo trafficking^7^. During aging, tau becomes hyperphosphorylated, resulting in misfolding and a decreased affinity for microtubules^8^. Tau pathology is transmitted cell-to-cell^2,9^, and the spread and levels of pathological tau strongly correlates with the degree of cognitive decline in AD patients^10^. Tau spread occurs through synaptically connected neuronal circuits; starting in the entorhinal cortex but eventually spreading to the hippocampus and neocortical areas^11^. Tau is released from neurons in an activity dependent fashion^12^ as free protein that is not enclosed in a membrane^13^ or in extracellular vesicles (EVs)^3–5^. Healthy neurons take-up extracellular free tau by mechanisms that include LRP mediated uptake^14^. However, EV tau uptake occurs by poorly delineated endocytic pathways^15^. When misfolded tau is taken up, it corrupts the conformation of physiological tau through a seeding process that ultimately leads to the formation of intracellular aggregates. Some of this newly misfolded tau is then released, possibly to protect cells from intracellular toxicity, continuing the cycle of cell-to-cell transmission. Released tau can be taken up by neurons or by microglia, which break down toxic proteins^16^. However, as the disease progresses, elimination and proteostasis pathways may become overloaded and contribute to the spread of tau pathology^16,17^. Interrupting the spread of tau pathology may be a promising therapeutic strategy for AD and other tauopathies, but the mechanisms of tau release and intercellular transmission remain poorly delineated.

Packaging of toxic proteins in EVs has been implicated in the pathology of various neurodegenerative disorders^18^. Exosomes, small (30-150nm) EVs usually derived from endosomal/multi-vesicular body compartments, may be involved in the initiation and propagation of AD pathology^16,19,20^. Endogenous tau protein is released from neurons in EVs^21^ and tau release is modulated by neuronal activity^12,22^. However, the exact type of EV, whether exosome or ectosomes^23^ (EVs released directly from the plasma membrane that can also be ∼100nm), critical for tau release is still unclear. EVs isolated from AD patient brains or transgenic mice expressing mutant human tau contain phosphorylated misfolded tau in EVs^4,5,24–26^. These brain-derived EVs can seed tau aggregation in cultured cells and in mouse brains^4,15,24,25,27^. EV-tau may be more potent at seeding tau aggregation, as transmission is more efficient than free tau protein^4^, and interfering with EV release reduces tau pathology in mice^4,16,28^. Nonetheless, the precise role of EV-tau and free tau in intercellular transmission of tau pathology remains to be determined. Understanding the balance and regulation of these two mechanisms of release has implications for therapeutic interventions using anti-tau antibodies^29^ which may not have access to membrane-encapsulated EV-tau.

The neuronal gene *Arc*, a master regulator of synaptic plasticity and memory consolidation^30^, has been implicated in AD pathology and synaptic dysfunction^31, 32^. Variations in the *Arc* gene in humans modulates risk for AD^33,34^ and Arc expression is disrupted in AD mouse models. Increased seizure activity and dysregulated Arc protein levels have also been observed in AD patients^32,35^. Arc protein spontaneously forms virus-like capsids, a property derived from its retrotransposon origins, that encapsulate RNA and are released in EVs that mediate intercellular transmission of proteins and RNAs^36^. The Drosophila Arc (*dArc*) homologs, which originated independently from distinct lineages of Ty3 retrotransposons, also form virus-like capsids^36,37^ that are packaged in EVs that transmit RNA from the neuromuscular junction synapse into muscle cells^38^. dArc1 protein levels, including high-molecular-weight species indicative of capsids, are increased in a tauopathy *Drosophila* model (tau^R406W^) and dArc1 modulates tau-induced neurodegeneration^39^. Endogenous EV-tau release is modulated by neuronal activity and high neuronal activity exacerbates the spread of tau pathology^12,21^. Hyper-phosphorylated tau is mislocalized in dendrites/post synaptic compartments^40–42^. Arc EV release is also enhanced by neuronal activity, and occurs in dendrites directly from the cell surface through the coordinated trafficking and assembly of Arc capsids via the I-BAR protein IRSp53^43^, which was recently found to also interact with tau^44^. Together, these data suggest that Arc may play an important role in regulating tau pathology. Here, we show that Arc plays a critical role in intercellular transmission of tau by packaging bioactive tau into neuronal EVs.

## Results

### Arc is critical for the release of tau in brain-derived EVs

Arc and tau are both released in EVs from neurons in response to neuronal activity^21,28,43^. Thus, we hypothesized that Arc may play a role in packaging tau into EVs. To test this *in vivo*, we isolated EVs from mouse brain tissue using ultracentrifugation followed by size exclusion chromatography (SEC) (Fig. S1A). This protocol circumvents homogenization of cells by gross chopping of the brain using a razor blade and enzymatic digestion of brain tissue, which prevents intracellular contaminants released due to the shear force applied during homogenization. Brain derived EVs were positive for common EV markers^45^, such as tetraspanins (CD9), Syntenin, and Alix, but were negative for the nuclear histone protein H3 (Fig. S1B). Nanoparticle tracking showed an enrichment for small EVs with an average size of 130nm and negative-stain transmission electron microscopy images showed similar sized vesicles (Fig. S1C). Using this protocol, we isolated brain EVs from 4-month-old rTg4510 (rTg^WT^) mice. These mice express multiple copies of the mutant (P301L) human tau (hTau) transgene in forebrain excitatory neurons under the CaMKII promoter^46^, and develop tau pathology relatively early (∼4-6 months). To identify hTau species packaged in EVs, we immunoblotted isolated brain EVs using human tau antibodies (Tau 13– tau amino acids 2-18^47^; Tau 22– putative oligomeric tau^48^; AT8 – phosphorylated 202/205 tau^49^; Tau Y9– phosphorylated tau at tyrosine 18^50^; Tau HT7 – tau amino acids aa 15-163^51^) that detect hTau (Fig. S2A). We detected hTau in brain derived EVs, using multiple tau antibodies (Fig. 1A), showing that brain-derived EVs contain full-length (molecular weight of ∼55kDa, which corresponds to the 0N4R isoform expressed in rTg4510 mice) and phosphorylated forms of hTau.

**Figure 1.**
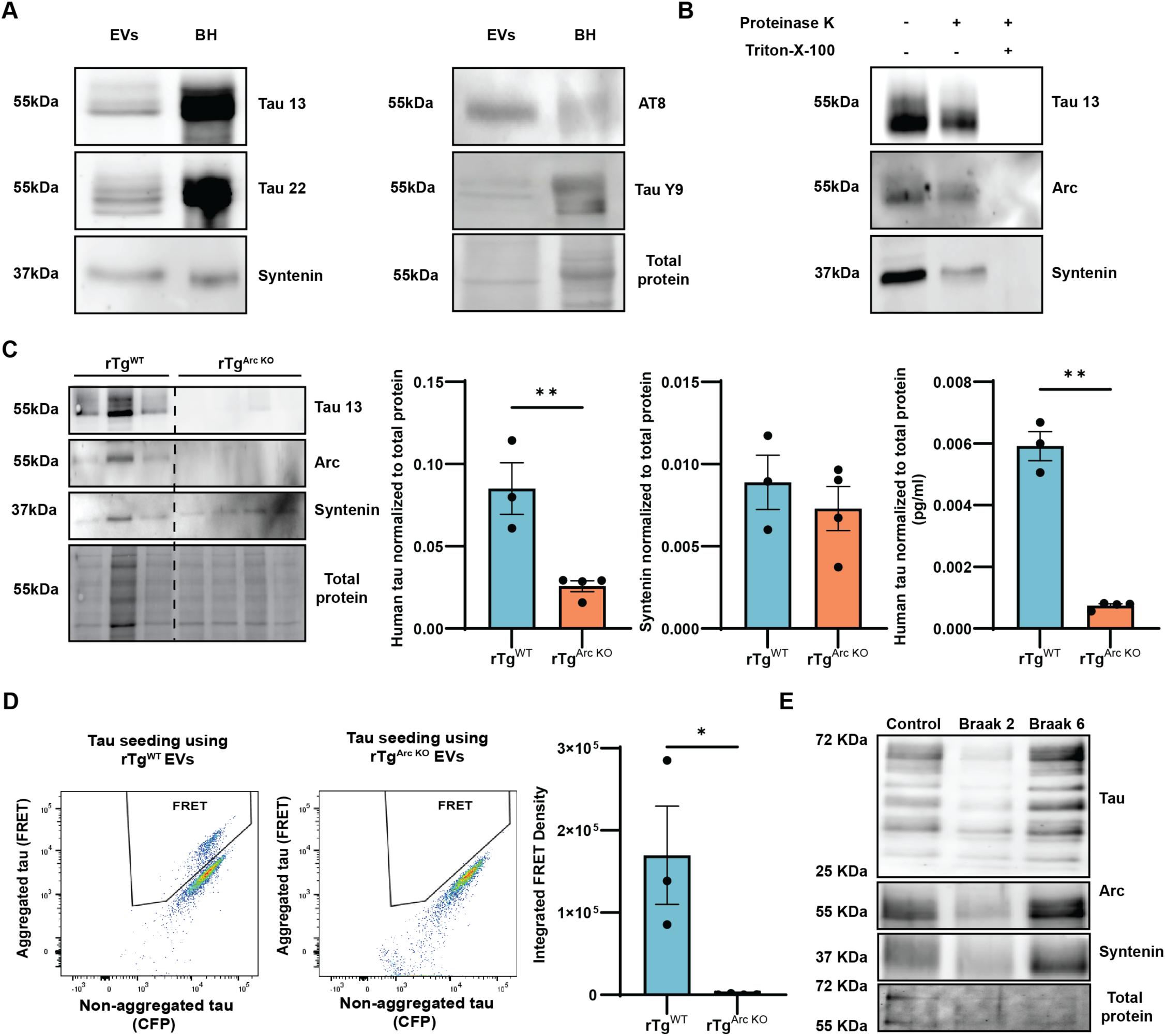
Arc is critical for the release of seed-competent hTau in EVs. **A.** *hTau is released in brain EVs isolated from rTg^WT^ mice.* EVs were isolated from brains of 4-month-old rTg^WT^ mice (n=3M). EVs and brain homogenates (BH) from 4-month-old rTg^WT^ mice were immunoblotted for hTau (Tau 13, Tau 22, AT8, and Tau Y9 antibodies) and Syntenin. Brain derived EVs contain hTau, including putative oligomeric forms as detected by Tau 13 and Tau 22 antibodies. **B.** *hTau and Arc in EVs is protected from Proteinase K degradation.* EVs were isolated from the brains of 4-month-old rTg^WT^ mice (n=3F). Fractions 1-4 obtained after SEC were pooled and incubated with Proteinase K (7ug/ml) with or without detergent (1% triton-X-100) for 10 mins. A representative western blot shows Arc, hTau, and Syntenin are protected from Proteinase K degradation when no detergent is present. **C.** *Arc facilitates the release of hTau in EVs isolated from rTg mice.* EVs were isolated from the brains of 4-month rTg^WT^ mice (n=6, 2M, 4F with each EV prep consisting of two mice) and rTg^Arc^ ^KO^ (n=8, 4M, 4F with each EV prep consisting of two mice). Fractions 1-4 were pooled and blotted for hTau (Tau 13), Arc, and Syntenin. (Dotted line indicates spliced blot to crop out irrelevant lanes). Levels of hTau were significantly reduced in the absence of Arc, while there was no difference in general EV production, as indicated by similar Syntenin levels. hTau levels in EVs isolated from rTg^Arc^ ^KO^ mice, as assessed by total hTau ELISA, are significantly reduced. **D.** *EV-hTau is capable of tau seeding.* The same EV samples used for hTau analysis were used in a FRET tau seeding assay. Equal amounts of rTg^WT^ and rTg^ArcKO^ EVs were transfected in HEK biosensor cells and FRET positive cells were counted using flow cytometry. FRET positive HEK biosensor cells exhibiting hTau aggregation are observed in the FRET gate in the representative flow cytometry plots. There are fewer FRET positive cells that were transfected with rTg^Arc^ ^KO^ EVs as compared to FRET positive cells transfected with rTg^WT^ EVs. The integrated FRET intensity (calculated by multiplying the percentage of FRET positive cells and median fluorescence intensity of FRET positive cells) of cells transfected with rTg^Arc^ ^KO^ EVs is significantly lower than cells transfected with rTg^WT^ EVs, indicating a lack of tau seeding. **E.** *Arc and hTau are present in EVs isolated from post-mortem human brain tissue.* EVs were isolated from a control, Braak stage 2, and Braak stage 6 post-mortem human brain tissue (Brodmann area 8/9). Isolated EVs were immunoblotted for Arc, hTau (Tau 13 antibody), and Syntenin. Arc and hTau are expressed in human brain-derived EVs. (Statistical analysis: Unpaired t-test, \**p*<0.05, \*\**p*<0.01).

**Supplementary Figure 1.**
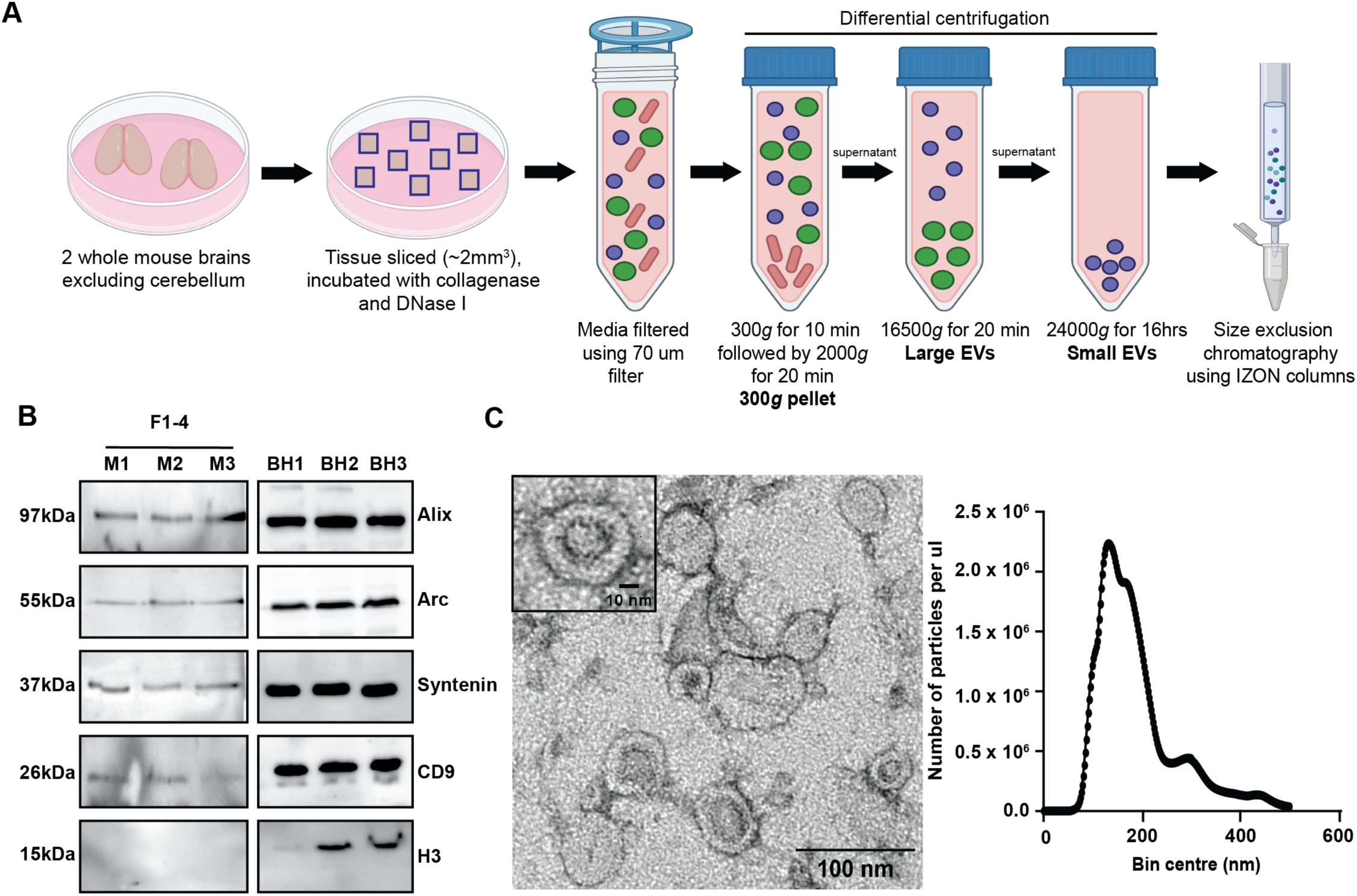
Characterization of small EVs isolated from mouse brain. **A.** Schematic overview of the brain EV isolation protocol. Flash-frozen mouse whole brains (excluding cerebellum) are sliced, collagenase and DNase treated, subjected to differential centrifugation, and then run through size exclusion chromatography using IZON columns. The early fractions (1-4) are enriched in small EVs (<200nm) and were used for further analysis. **B**. Early SEC fractions (F1-4) from 3-month-old WT mouse brains were pooled and concentrated. Three brain EV preparations that consisted of two mouse brains each (M1, M2, M3) and brain homogenates (BH1, BH2, BH3) were stained for positive EV markers (Alix, CD9, Syntenin), a negative EV marker (H3), and Arc. **C.** Representative negative-stain transmission electron microscopy image and size distribution nanoparticle tracking plots show an enrichment of small EVs with an average size of 100-130nm.

**Supplementary Figure 2.**
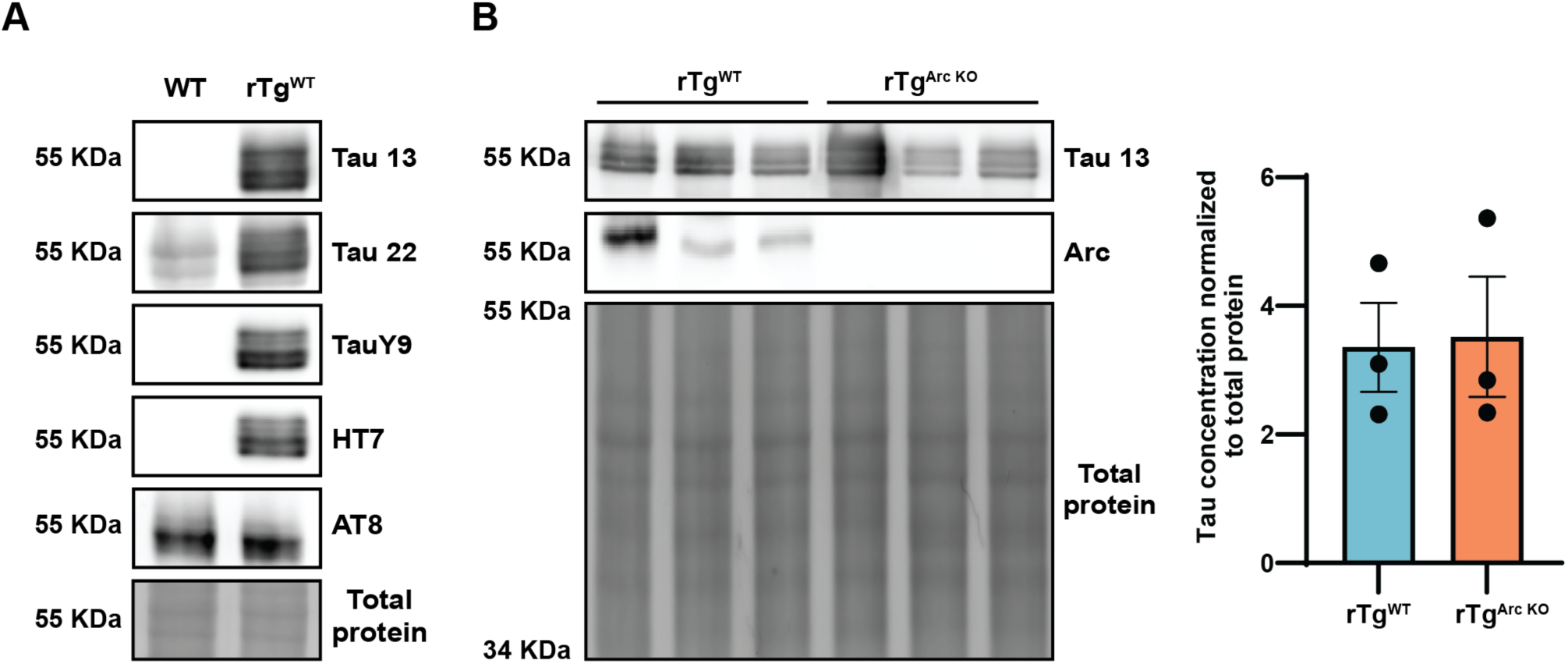
rTg^WT^ and rTg^Arc^ ^KO^ mice have comparable total levels of hTau at 4 months of age. **A.** *Specificity of tau antibodies for hTau.* Brain homogenates were collected from 4-month-old WT and rTg^WT^ mice. The brain homogenates were blotted for tau using Tau 13, Tau 22, Tau Y9, HT7, and AT8 antibodies. Tau 13, Tau Y9, and HT7 specifically detect hTau and not endogenous mouse tau. **B.** *hTau is expressed in comparable amounts in 4-month-old rTg^WT^ and rTg^Arc^ ^KO^ mice.* Brain homogenates were collected from 4-month-old rTg^WT^ and rTg^Arc^ ^KO^ mice (n=3, 1M, 2F). Levels of hTau were analyzed by western blot using Tau 13 antibody. rTg^WT^ and rTg^Arc^ ^KO^ mice have comparable amounts of total hTau.

We detected Arc, hTau, and Syntenin in brain-derived EVs isolated from 4-month-old rTg^WT^ mice (Fig. 1B). To confirm that hTau and Arc are inside EVs, we conducted a proteinase K protection assay that showed Arc and hTau are protected from proteinase K degradation (Fig. 1B) in the absence of detergent. To determine if Arc facilitates the release of hTau in brain EVs, we crossed rTg^WT^ mice with homozygous germline Arc knock-out mice (rTg^Arc^ ^KO^). rTg^WT^ and rTg^Arc^ ^KO^ mice express comparable amounts of hTau (Fig. S2B). hTau levels in EVs extracted from 4-month-old rTg^WT^ and rTg^Arc^ ^KO^ were assessed by western blot and total hTau ELISA. hTau levels in EVs isolated from rTg^Arc^ ^KO^ are dramatically decreased, while Syntenin levels are comparable to rTg^WT^ mice (Fig. 1C). This suggests Arc is critical for packaging hTau into brain EVs.

### Brain-derived EV-hTau is capable of seeding tau pathology

Neurons take up extracellular misfolded tau, which alters the conformation of physiological tau to seed the formation of tau aggregates^2^. EVs isolated from AD patients are capable of seeding tau pathology^4^. Brain EVs isolated from rTg^WT^ have higher tau levels compared with EVs isolated from rTg^Arc^ ^KO^ mice, suggesting that Arc may also regulate tau seeding potential. To determine tau seeding potential, we conducted a FRET-based tau seeding assay in HEK293 biosensor cells^52^. Tau biosensor HEK cells stably express hTau(P301S)^RD^-CFP/YFP.

The hTau P301S repeat domain (RD) does not aggregate on its own, but aggregates in the presence of tau seeds, generating a FRET signal. We transfected 10μg of rTg^WT^ or rTg^Arc^ ^KO^ brain EVs in HEK biosensor cells expressing hTau(P301S)^RD^-CFP/YFP using lipofectamine and quantified the number of FRET positive cells by flow cytometry. rTg^WT^ brain derived EVs induced FRET positive cells (Fig. 1D). In contrast, rTg^Arc^ ^KO^ EVs were unable to induce FRET positive cells (Fig. 1D). The integrated FRET density, which represents the total tau seeding of the transfected EVs, was significantly reduced in rTg^Arc^ ^KO^ EV transfected tau biosensor HEK cells (Fig. 1D). These observations show that rTg^WT^ brain EVs are capable of seeding tau pathology, whereas rTg^Arc^ ^KO^ brain EVs do not have significant tau seeding potential, most likely due to the low levels of hTau in these EVs.

### Human brain-derived EVs contain Arc and hTau

Mouse brain-derived EVs contain Arc and hTau. However, mutant hTau is over-expressed in this mouse line. To determine whether human brain-derived EVs contain Arc and hTau, we obtained flash-frozen postmortem Brodmann area 8/9 prefrontal cortex tissue from a healthy control, a Braak stage 2 AD patient, and a Braak stage 6 AD patient (see methods for patient details). We isolated EVs from flash-frozen post-mortem human brain tissue using a previously described protocol^4^ (see methods). We detected the EV marker Syntenin (Fig. 1E), confirming successful isolation of human EVs. We also detected hTau and Arc in human brain-derived EVs (Fig. 1E). Interestingly, brain tissue obtained from the late stage (Braak 6) AD patient has higher levels of Arc, hTau, and Syntenin. These data suggest that Arc may regulate the release of EV-tau in human brains and corroborates the observations of Arc and EV-tau in rTg mice.

### Arc mediates the release of hTau in Arc-IRSp53 EVs

Tau is released from neurons as free tau or in membrane enveloped EVs. To directly test if Arc facilitates the release of free or EV-hTau from neurons, we transduced WT and Arc KO primary cortical neuron cultures with hSyn-eGFP-2A-human tau (P301L) on days in vitro (DIV) 7. Cell lysates and media were collected from transduced neurons at DIV 18, 48 hours post a full media change. We measured total hTau levels using a human tau specific ELISA with or without 1% triton-X-100 detergent. The non-detergent sample only detects free hTau, while the detergent sample lyses EVs, allowing quantification of total hTau levels and thus the protected/EV hTau fraction. The release of free hTau is comparable between WT and Arc KO neuronal cultures (Fig. 2A). However, the release of EV-hTau is significantly reduced in Arc KO primary neurons (Fig. 2A).

**Figure 2.**
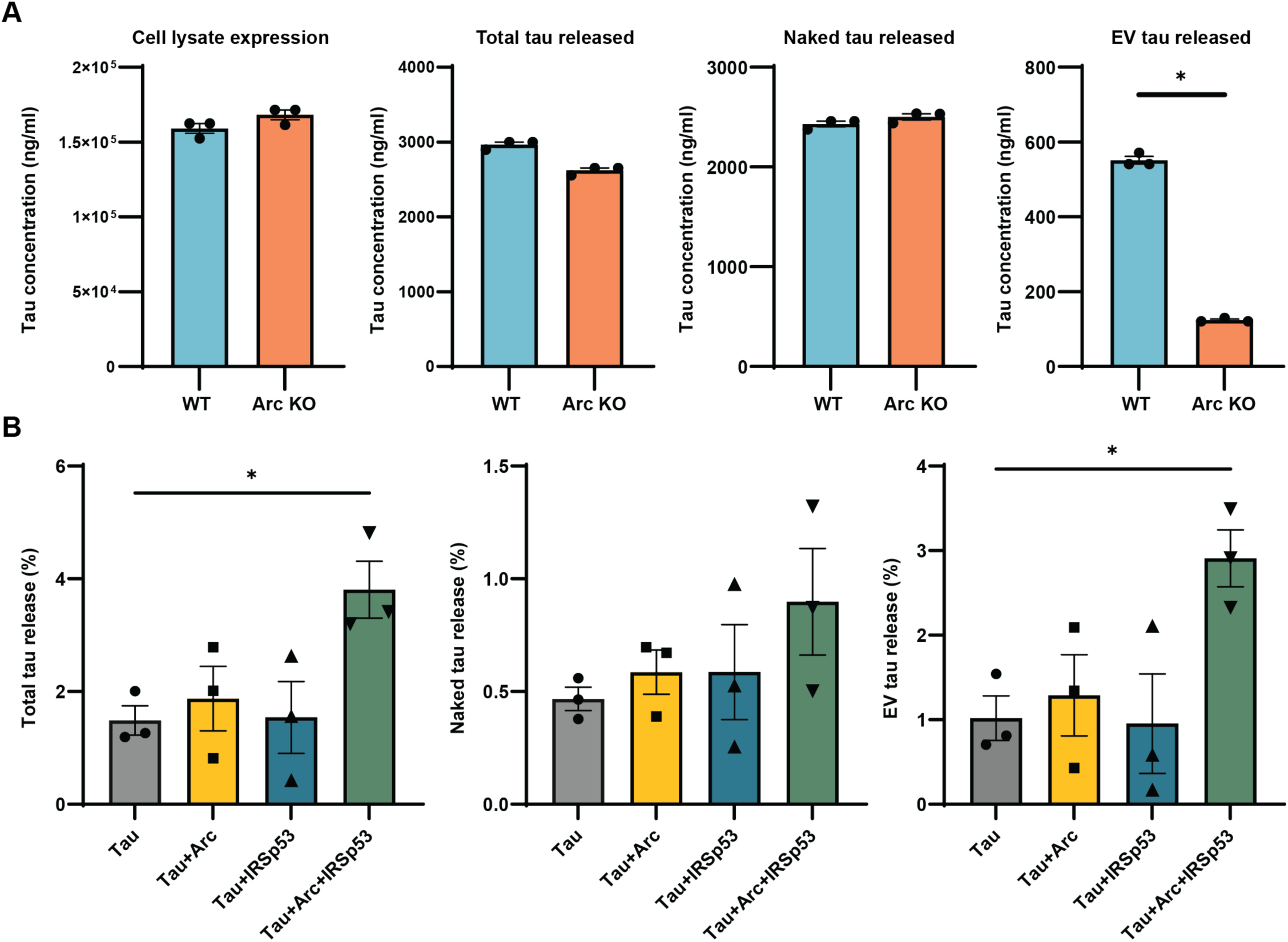
hTau is preferentially released in Arc-IRSp53 EVs. **A.** WT and Arc KO primary cortical neurons were transduced with hSyn-eGFP-2A-hTau*P301L at DIV 7 and media was collected on DIV 18. hTau levels in the media were quantified with or without detergent (1% triton-X-100) using a hTau ELISA to quantify total hTau and naked hTau, respectively. A significant decrease in EV-hTau levels was observed in Arc KO neurons, while naked and total hTau release were similar to WT neurons. (Statistical analysis: Unpaired t-test, *p<0.05 – data points are technical replicates). **B.** *hTau is released in Arc-IRSp53 EVs.* Neuro2A cells were transfected with P301L hTau-Hibit alone or with Arc and/or IRSp53 for 24 hours. Conditioned media was collected 24 hours post full media change. Luminescence was measured with or without detergent to quantify total hTau or free hTau in the media. Free hTau release was not affected by Arc and/or IRSp53 expression. However, EV-hTau release was significantly enhanced when Arc and IRSp53 were co-expressed. (Statistical analysis: One-way ANOVA, *p<0.05 – data points are technical replicates).

It is possible that the phenotypes observed in Arc KO neurons could be due to indirect mechanisms that result from loss of Arc protein. For example, hTau release could be affected by neuronal activity levels. Previous studies showed that Arc KO primary neurons have increased surface AMPARs and larger mini-EPSC amplitudes^53^, although *in vivo* basal synaptic transmission is normal in the hippocampus of Arc KO mice^54^. We find that primary Arc KO neurons have similar spontaneous neuronal activity and intrinsic excitability as WT neurons (Fig. S3). However, previous studies have shown that high neuronal activity leads to an *increase* in tau release ^12,22^, thus it seems unlikely that increased neuronal activity in Arc KO mice would result in *less* tau release.

Arc EV biogenesis and release is mediated by the I-BAR protein IRSp53^43^. To test whether EV-hTau release is regulated by IRSp53, we used a split-nano luciferase system^55^. We cloned the small HiBiT tag (11 amino acids) onto the C-terminus of P301L hTau and transfected Neuro2A cells with hTau-HiBiT alone or hTau-HiBiT+Arc with or without IRSp53. Cell lysate and media samples were collected to measure luciferase luminescence, which occurs only when the HiBiT tag binds Large-bit which is added to the media. Total hTau and EV-hTau release is significantly enhanced when both IRSp53 and Arc are expressed (Fig. 2B), but not when only Arc or IRSp53 is expressed. These data show that hTau is specifically packaged into Arc EVs generated by IRSp53.

**Supplementary Figure 3.**
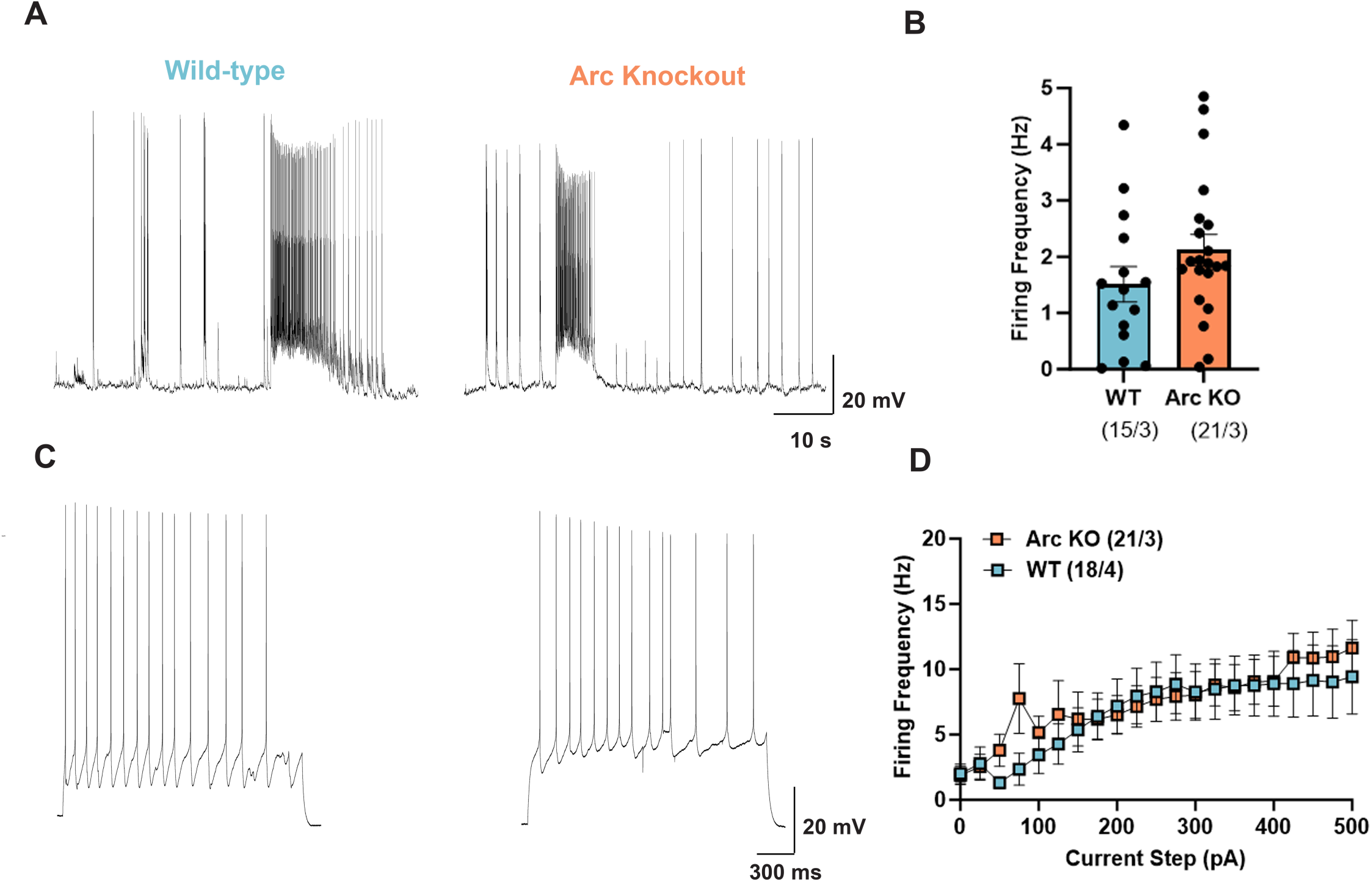
Arc KO primary hippocampal neurons have comparable spontaneous neuronal activity and excitability to WT neurons. **A.** Representative recordings of spontaneous neuronal firing of DIV 16 WT and Arc KO primary cultured hippocampal neurons. **B.** There is no significant difference in the average spontaneous firing frequency between WT neurons (n=15 from 3 cultures) and Arc KO neurons (n=21 from 3 cultures). **C.** Representative recordings of DIV 16 WT and Arc KO hippocampal neurons in response to 300 pA current injection. **D.** WT (n=18 from 4 cultures) and Arc KO hippocampal neurons (n=21 from 3 cultures) have comparable firing frequency in response to current injections.

### Arc, IRSp53, and hTau interact in mouse and human cortex

Arc may package hTau into EVs through a direct protein interaction. To determine if hTau binds Arc, we conducted GST-pulldown experiments. We purified GST, GST-Arc, or GST-Endophilin from *E. coli* as described previously^36^. We incubated immobilized purified protein on beads and incubated them with HEK293 cell lysate from untransfected cells (UT), or cells transfected with WT or P301L hTau. WT and P301L hTau binds GST-Arc, but not GST or GST-Endophilin (Fig. 3A). Purified recombinant mutant hTau protein also binds purified Arc protein (Fig. 3B), indicating a direct interaction. Since Arc EV release is mediated by IRSp53^43^, we determined whether Arc, hTau, and IRSp53 interact *in vivo*. We conducted immunoprecipitation experiments from mouse cortex. To induce Arc expression, 4-month-old rTg^WT^ mice were placed in an enriched environment (toys and novel objects in the home cage) for 6 hours. Arc was immunoprecipitated from cortical lysates and the elutes were blotted for Arc, hTau, and IRSp53. Both hTau and IRSp53 co-immunoprecipitated with Arc (Fig. 3C), showing that Arc-hTau-IRSp53 interact in a complex in the mouse brain *in vivo*.

**Figure 3.**
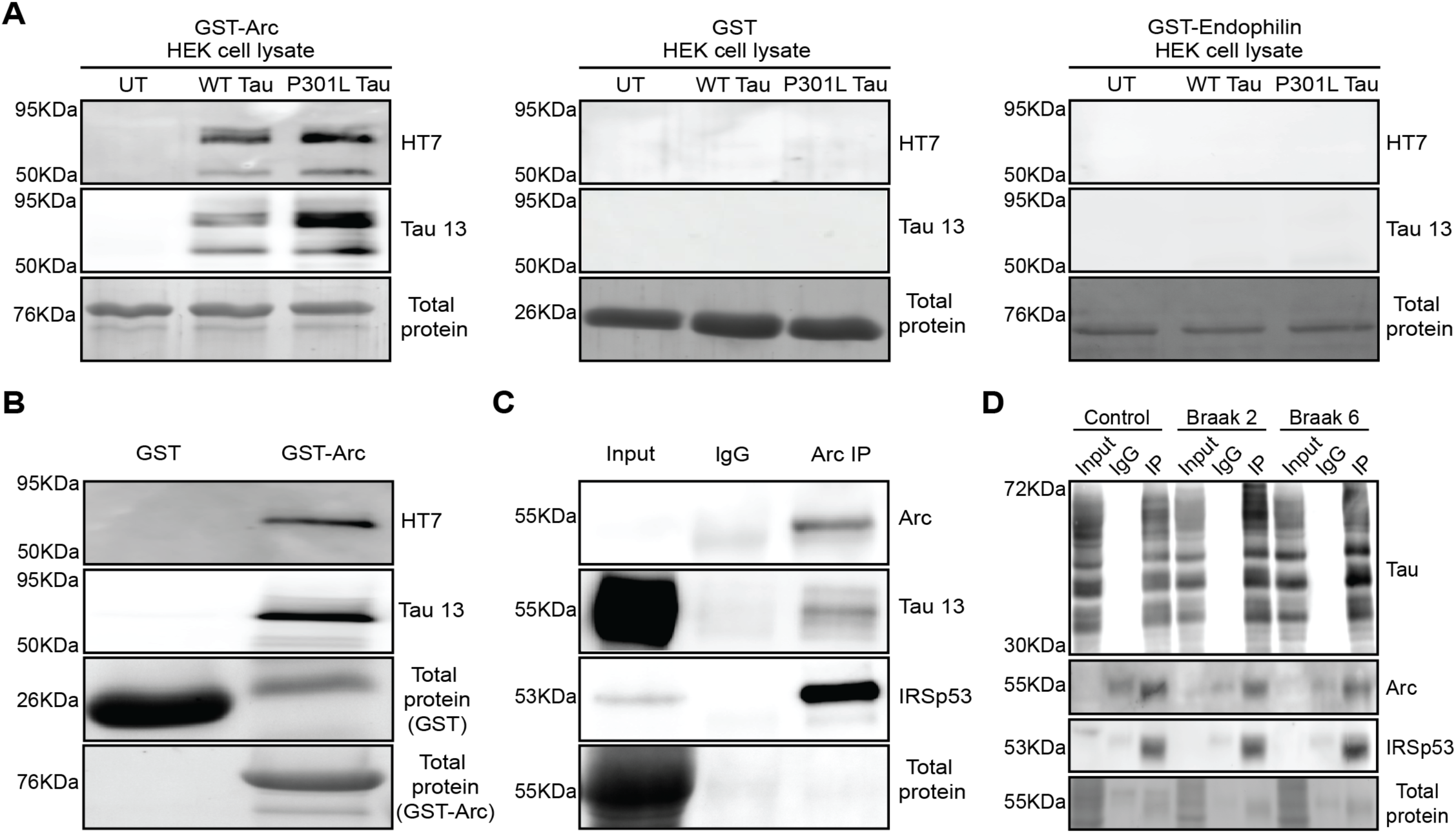
Arc directly binds to hTau and forms a complex with IRSp53 in mouse and human cortex. **A.** *hTau binds Arc.* GST-Arc, GST, or GST-Endophilin were immobilized on beads and incubated with HEK cell lysate from untransfected (UT), WT hTau (2N4R) transfected, or mutant (P301L) hTau (2N4R) transfected cells. hTau was not present in the eluted fractions collected from GST or GST-Endophilin but was detected in fractions collected from GST-Arc. **B.** *Arc directly binds hTau.* GST or GST-Arc was immobilized on beads and incubated with purified recombinant hTau (P301L). hTau was present in the eluted fraction collected from GST-Arc but not GST. **C. h***Tau co-immunoprecipitates with Arc and IRSp53 in mouse cortex.* Cortical tissue was collected from 4-month-old rTg^WT^ mice that experienced 6 hours of enrichment to increase Arc expression. Brain homogenates were incubated with IgG (rabbit polyclonal, EMD Millipore) or Arc antibody (rabbit polyclonal, Synaptic systems). Input and IP samples were blotted for Arc (mouse monoclonal, Santa Cruz), hTau (mouse, Tau13 antibody), and IRSp53 (mouse, Abcepta). hTau was detected in Arc-IPs but not in IgG-IPs. **D.** *Arc and IRSp53 co-immunoprecipitates with hTau in human post-mortem prefrontal cortical tissue*. Human post-mortem brain tissue from control, Braak stage 2, and Braak stage 6 AD patients were incubated with IgG (rabbit mouse antibody, Cell signaling) or D5D8N hTau antibody (rabbit mouse antibody, Cell signaling). Input and IP samples were blotted for hTau (mouse monoclonal, Tau 13 antibody, Biolegend), Arc (purified E5 alpaca recombinant nanobody), and IRSp53 (mouse monoclonal, Abcepta). Arc and IRSp53 were specifically detected in hTau-IPs.

To determine whether Arc and IRSp53 interact with hTau in human brain, we immunoprecipitated hTau from post-mortem cortical tissue of a healthy control, a Braak stage 2 AD patient, and a Braak stage 6 AD patient (the same samples used for EV isolations). Both Arc and IRSp53 co-immunoprecipitated with hTau in control and AD patient brains (Fig. 3D). Together, these data show that Arc and IRSp53 form a complex with hTau in both mouse and human cortex.

### Molecular modelling of Tau-Arc interactions suggests the formation of a fuzzy complex

To gain insights into how hTau interacts with Arc, we conducted all-atom molecular dynamics simulations using the full-length human and mouse Arc with the four repeats of the hTau 2N4R (hTau, residues 244-369). The simulations indicate that hTau remains largely disordered in complex with both human and mouse Arc, forming multisite, dynamic interactions (Fig. S4A-C). The interactions with the N-terminal coiled-coiled region appear to be more stable than those with the Gag domain, while contacts with the C-terminal region are transient (Fig. S4C-D). Despite the lack of regular secondary structures, the QIVY motif of hTau (residues 307-310), which is critical for Tau seeding^56,57^, adopts a β-strand conformation (Fig. S4A, inset) that interacts with Arc (Fig. S4E). This local sequence element contacts both the Arc coiled-coil region and the N-terminal part of the Gag domain (Fig S4C-D) and can form a parallel β-sheet with the Arc motif TQIF (residues 211-214) (Fig. S4A, inset, S4B). The mutant P301L hTau expressed in rTg mice appears to form more stable contacts with mouse Arc as compared to WT hTau, in particular through the PHF6 and PHF6* motifs, which are aggregation prone regions^56,57^ (Fig. S4D,). In addition, P301L hTau exhibits more stable β conformations (Fig. S4G) than WT hTau while in a complex with Arc. The complex shows mixed anti-parallel hTau β-sheets with short β-elements of mouse Arc (Fig. S4F). These data suggest that hTau and Arc form a fuzzy assembly^58^ that has dynamic and variable contact patterns, which may mask hTau aggregation-prone elements to limit hTau self-assembly.

**Supplementary Figure 4.**
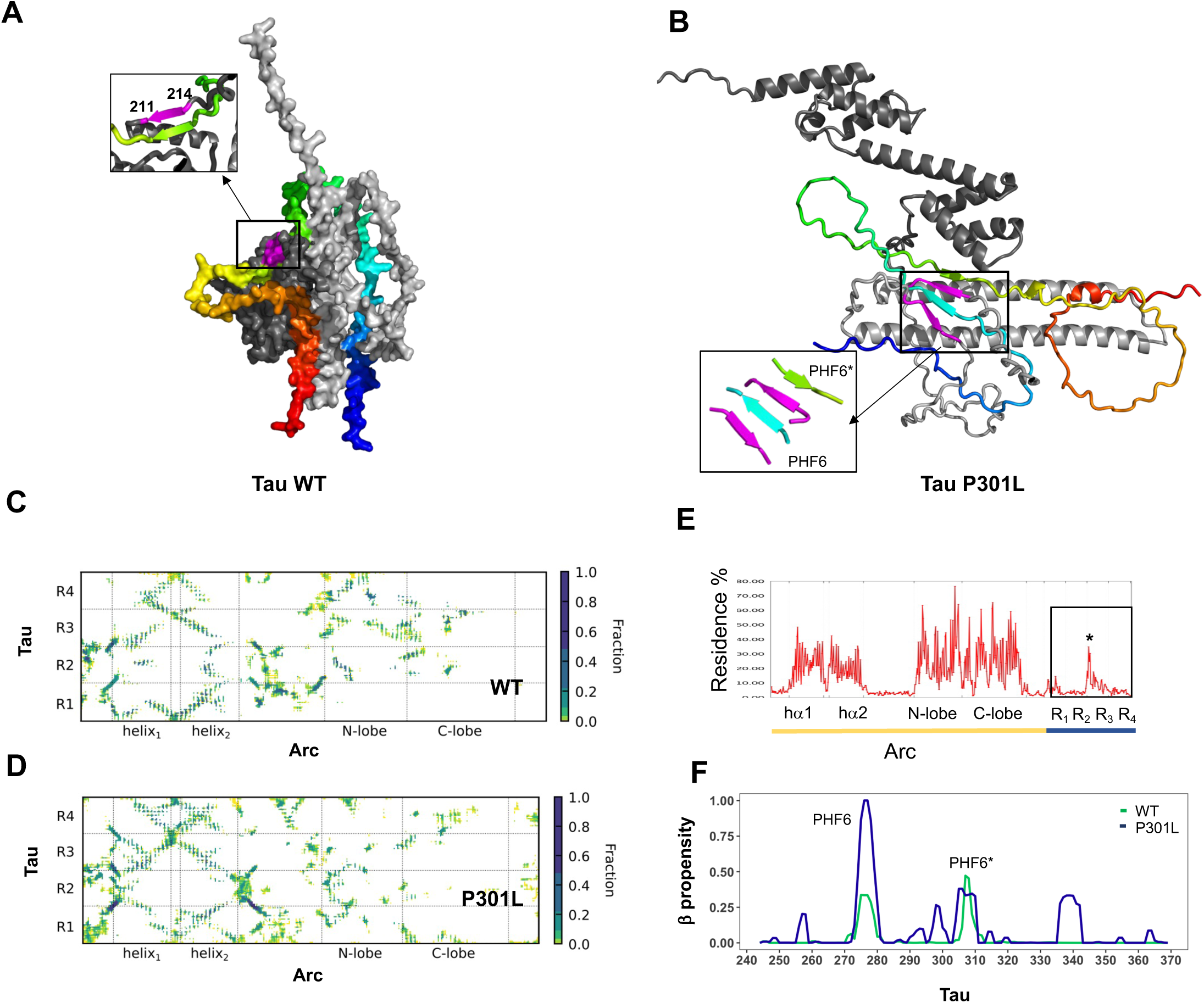
Tau forms a fuzzy complex with Arc through multisite, dynamic interactions. **A.** *Representative surface model of hTau and Arc interactions*. Tau repeats R_1_-R_4_ (UniProt code P10636-8; colored residues 244-369) interact with both the N-terminal coiled-coil (light gray) as well as the Gag domain (dark gray). Inset: Tau QIVY motif (residues 307-310 in 2N4R) form a β-strand (lime), which parallels a short β-structure in Arc (residues 211-214, UniProt code Q7LC44, magenta). **B***. Representative structure of P301L hTau with Arc*. hTau R_1_-R_4_ repeats (colored ribbons) establish multisite contacts with Arc (gray), P301L hTau forms a mixed β-sheet with mouse Arc (inset). **C-D.** *Contact maps of hTau-Arc interactions*. hTau repeats R_1_-R_4_ (UniProt code P10636-8) interact with both the N-terminal coiled-coil as well as the Gag domain of mouse Arc (UniProt code Q9WV31). The P301L hTau mutant (***D***) exhibits more stable contacts with mouse Arc than WT hTau (***C***), especially with the R_2_-R_3_ repeats. The persistence of contacts is shown as the fraction of snapshots, using a threshold ≤ 6 Å between the closest heavy atoms. **E.** *Persistence of Tau β-element interactions with Arc*. Residue interaction residence times (both intra- and intermolecular) are shown as a percentage of the simulation time. The hTau β-structure (residues 307-310 in 2N4R marked by star) is engaged most frequently in self and Arc interactions. **F.** *β-structure propensity is increased in P301L mutant hTau*. P301L hTau (blue) exhibits more β-structures as compared to WT hTau (green) with increased stability for the PHF6* motif. The propensity of β-structures was computed with the STRIDE algorithm (see methods).

### Intracellular hTau accumulates in rTg^Arc^ ^KO^ neurons, which is correlated with accelerated cell loss in the hippocampus

Arc-dependent release of hTau in EVs may play a role in the elimination of intracellular misfolded hTau. To test this hypothesis, we collected brains from male and female rTg^WT^ and rTg^Arc^ ^KO^ littermates (n=8, 3M, 5F) at 4 months of age, the same age where we observed significantly less hTau in rTg^Arc^ ^KO^ EVs. We then assessed hTau levels in the CA1 region of the dorsal hippocampus. rTg^Arc^ ^KO^ mice have significantly increased intracellular hTau levels in individual neurons and dendrites, as compared with rTg^WT^ mice (Fig. 4A, C). However, there are also fewer neurons in the CA1 cell body layer of rTg^Arc^ ^KO^ mice (Fig. 4B, D). This suggests that hTau accumulates in neurons in the absence of Arc, which may ultimately cause higher levels of intracellular toxicity. We did not observe significant differences in the number or intensity of Iba1 positive microglia in rTg^Arc^ ^KO^ mice (Fig. S5), and no overt changes in morphology, suggesting that microglia are not altered in rTg^Arc^ ^KO^ mice.

**Figure 4.**
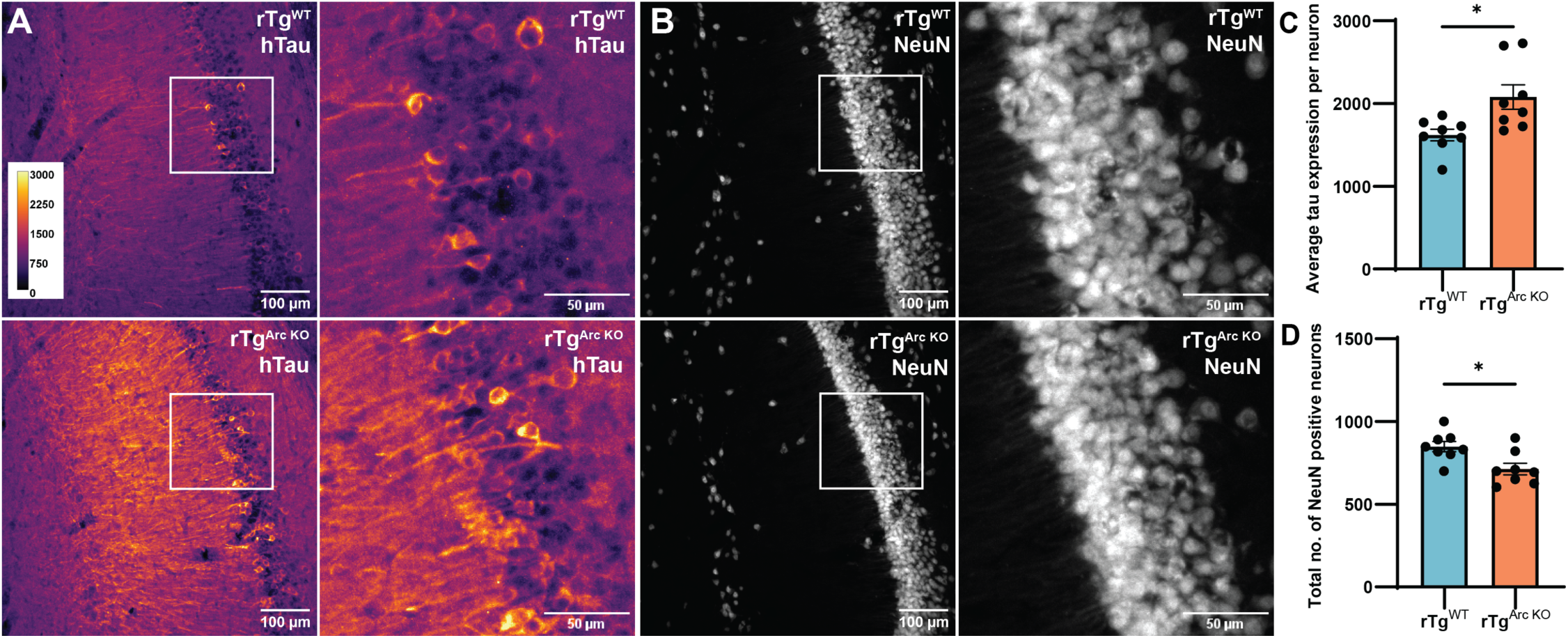
Intracellular hTau accumulates in neurons and there are less neurons in the cell-body layer of dorsal CA1 hippocampus in rTg^Arc^ ^KO^ mice. Brain slices were collected from 4-month-old rTg^WT^ and rTg^Arc^ ^KO^ littermates (n=8, 3M, 5F). Brain slices of the CA1 region of the dorsal hippocampus were immunostained for hTau (Tau13 antibody) and NeuN. **A.** Representative images of hTau staining in rTg^WT^ and rTg^Arc^ ^KO^ CA1. **B.** Representative images of NeuN staining in the same sections from A. **C.** Quantification of average hTau expression per neuron in the cell body layer. rTg^Arc^ ^KO^ mice have significantly higher intracellular hTau levels. **D.** Quantification of the total number of NeuN positive neurons in dorsal CA1. rTg^Arc^ ^KO^ mice have significantly lower NeuN positive cells in the CA1 cell body layer. (Statistical analysis: Unpaired t-test, \**p*<0.05).

hTau expression is very high in rTg mice and it is possible that the phenotypes we observed may be due to the transgene or mouse line. To further investigate whether Arc regulates intracellular levels of hTau, we crossed 3xTg AD (3xTg^WT^) mice with Arc KO mice to generate 3xTg^Arc^ ^KO^ mice. 3xTg AD mice express a mutant human APP (swe) and hTau (P301L) transgene driven by the *thy1* promoter, and a knock-in of mutant (M146V) PS1^59^. 3xTg AD mice develop amyloid plaques at 6 months and significant tau pathology by 12 months^59^. We evaluated tau pathology in 12-month-old 3xTg^WT^ and 3xTg^Arc^ ^KO^ mice (n=8, 3M, 5F). We imaged the CA1 region of the dorsal hippocampus and quantified hTau levels in the cell body and dendritic layer. As previously reported^60^, we observed sex differences at this age, with females exhibiting more severe tau pathology. There were no significant differences in overall hTau levels between 3xTg^WT^ and 3xTg^Arc^ ^KO^ male mice at this age (Fig. S6A-E), possibly due to low levels of tau pathology. However, 3xTg^Arc^ ^KO^ females have significantly higher hTau levels in the cell body and dendritic layers than 3xTg^WT^ females (Fig. S6F-H). 3xTg^Arc^ ^KO^ females also exhibit higher intracellular hTau levels in individual neurons, but fewer hTau positive neurons (Figure S6I-J). Together, these data indicate that intracellular hTau accumulates in neurons in the absence of Arc, which may ultimately exacerbate cell death, and is consistent with Arc-dependent release of hTau from neurons.

**Supplementary Figure 5.**
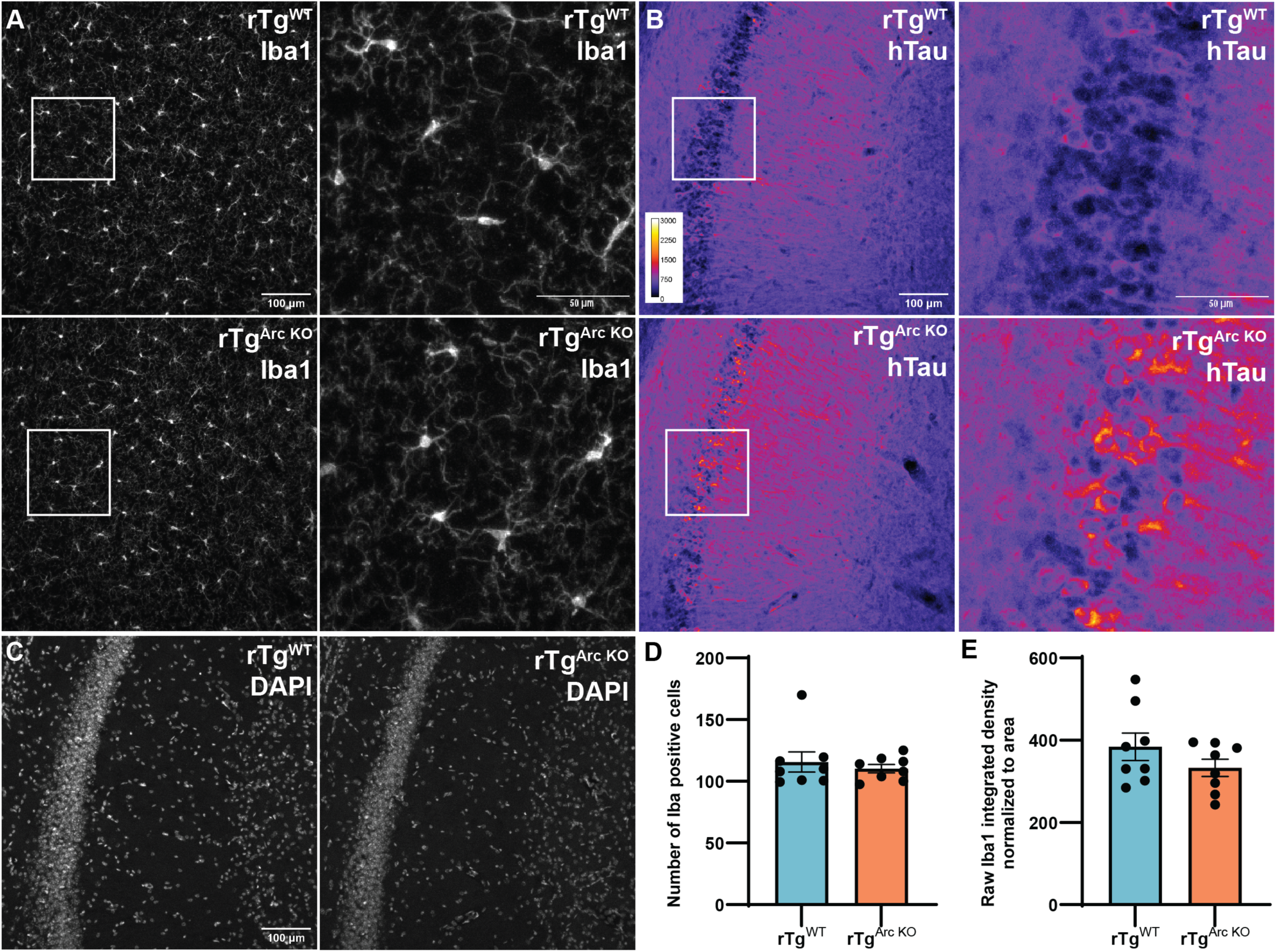
Microglia are comparable between 4-month-old rTg^WT^ and rTg^Arc^ ^KO^ littermate mice in dorsal CA1. Brain slices were collected from 4-month-old rTg^WT^ and rTg^Arc^ ^KO^ littermate mice (n=8, 3M, 5F). Brain slices were immunostained for Iba1, hTau (Tau13 antibody) and DAPI, and the CA1 region of the dorsal hippocampus was imaged. **A.** Representative images of Iba1 staining. **B.** Representative images of hTau staining. **C.** Representative images of DAPI staining. **D.** Quantification of the number of Iba1-positive cells. There are no significant differences between 4-month-old rTg^WT^ and rTg^Arc^ ^KO^ littermate mice. **E.** Quantification of raw Iba1 integrated density/area. There are no significant differences between old rTg^WT^ and rTg^Arc^ ^KO^ littermate mice. (Statistical analysis: Unpaired t-test).

**Supplementary Figure 6.**
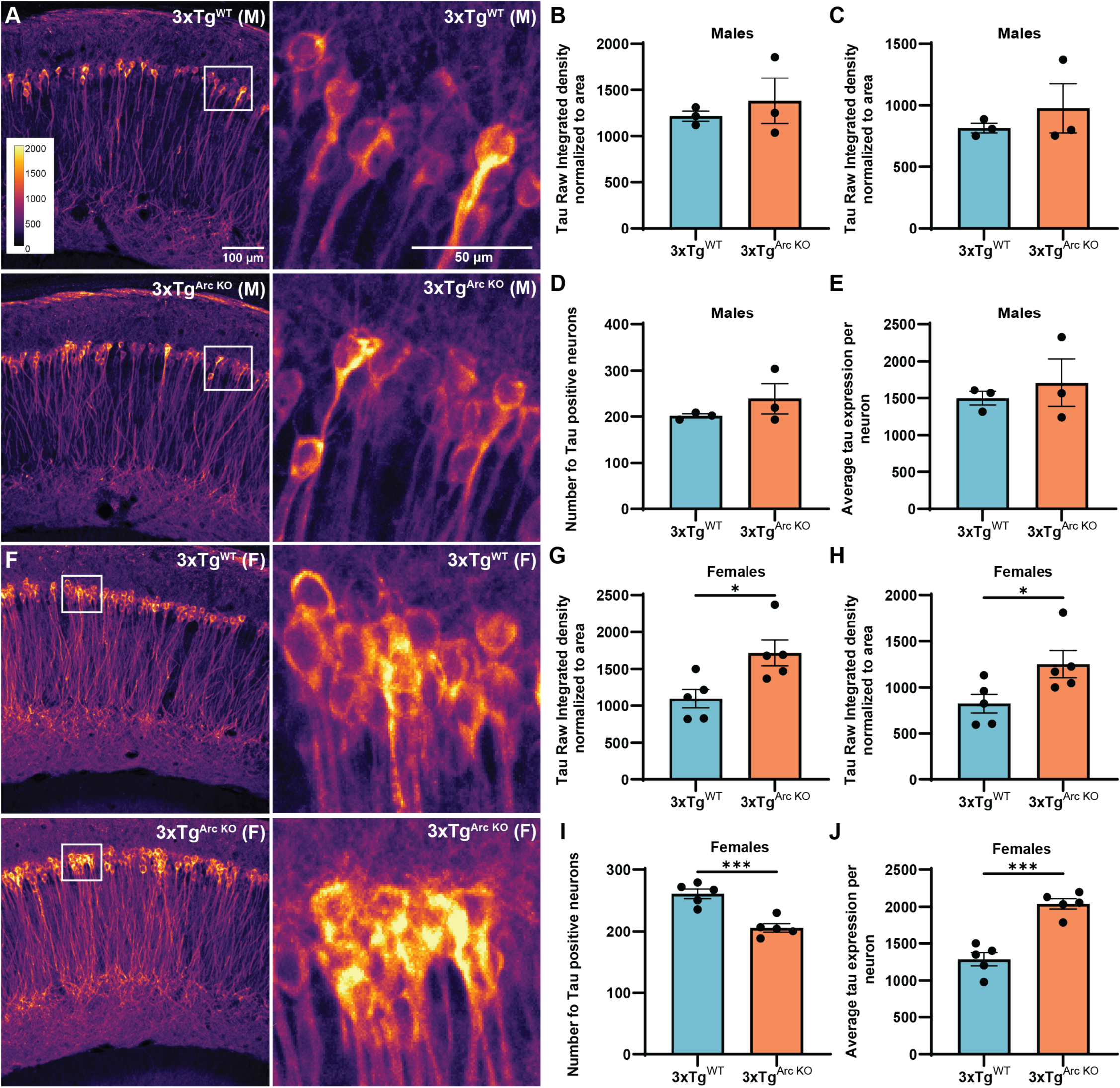
Intracellular hTau levels are increased in 3xTg^Arc^ ^KO^ female mice. 12-month-old 3xTg^WT^ and 3xTg^Arc^ ^KO^ brains (n=8, 3M, 5F) were collected and sectioned. Immunohistochemistry for hTau was performed using an Tau13 antibody. Imaging was performed in the CA1 region of the dorsal hippocampus. **A.** Representative images of hTau staining in 3xTg^WT^ and 3xTg^Arc^ ^KO^ males. **B.** Quantification of hTau levels in cell bodies normalized to area (male mice). **C.** Quantification of hTau levels in dendrites (male mice). **D.** Quantification of the number of hTau positive neurons (male mice). **E.** Quantification of the average hTau intensity per neuron (male mice). No significant differences were observed (**B-E**) in hTau levels in male mice. **F.** Representative images of hTau staining in 3xTg^WT^ and 3xTg^Arc^ ^KO^ female mice. **G.** Quantification of hTau levels in cell bodies normalized to area (female mice). **H.** Quantification of hTau levels in dendrites (female mice). **I.** Quantification of the number of hTau positive neurons (female mice). **J.** Quantification of the average hTau intensity per neuron (female mice). Intracellular hTau levels in neurons and dendrites was higher, while the number of hTau positive neurons was significantly reduced in 3xTg^Arc^ ^KO^ female mice. (Statistical analysis: Unpaired t-test, *p<0.05, ***p<0.001).

### Arc is critical for intercellular transmission of hTau

The transmission of tau is dependent on the efficiency of release and uptake^61^. While most tau is released in a free form, EV-tau may be taken up more efficiently^62^. Our data shows that Arc is critical for the release of EV-hTau, thus we hypothesized that Arc may regulate intercellular transmission of hTau. To determine whether Arc plays a role in intercellular hTau transmission, we employed a hTau transfer assay that uses an AAV construct, hSyn-eGFP-2A-hTau (P301L), which produces equimolar ratio of eGFP and hTau protein from a single mRNA^63^ (Fig. 6A). The 2A peptide sequence self-cleaves and separates eGFP and hTau during translation. As a result, transduced neurons express both eGFP and hTau as individual proteins in equal amounts. In contrast, neurons that receive hTau via intercellular transmission express hTau but not GFP. Thus, donor and recipient neurons can be identified by co-immunofluorescence of eGFP and hTau^63^ (Fig. 6A). To determine whether Arc is necessary for intercellular tau transmission *in vitro*, we sparsely transduced WT and Arc KO primary hippocampal neuron cultures at DIV 7 with AAV2/5: hSyn-eGFP-2A-hTau*P301L. 14 days post-transduction we stained for eGFP and hTau, and quantified the number of donor/recipient neurons. We found that the percentage of transduced donor neurons (GFP^+^, hTau^+^) is the same in WT and Arc KO neurons, but the percentage of hTau recipient neurons (GFP^−^, hTau^+^) is significantly reduced in Arc KO neurons (Fig. 6B-D).

**Figure 6.**
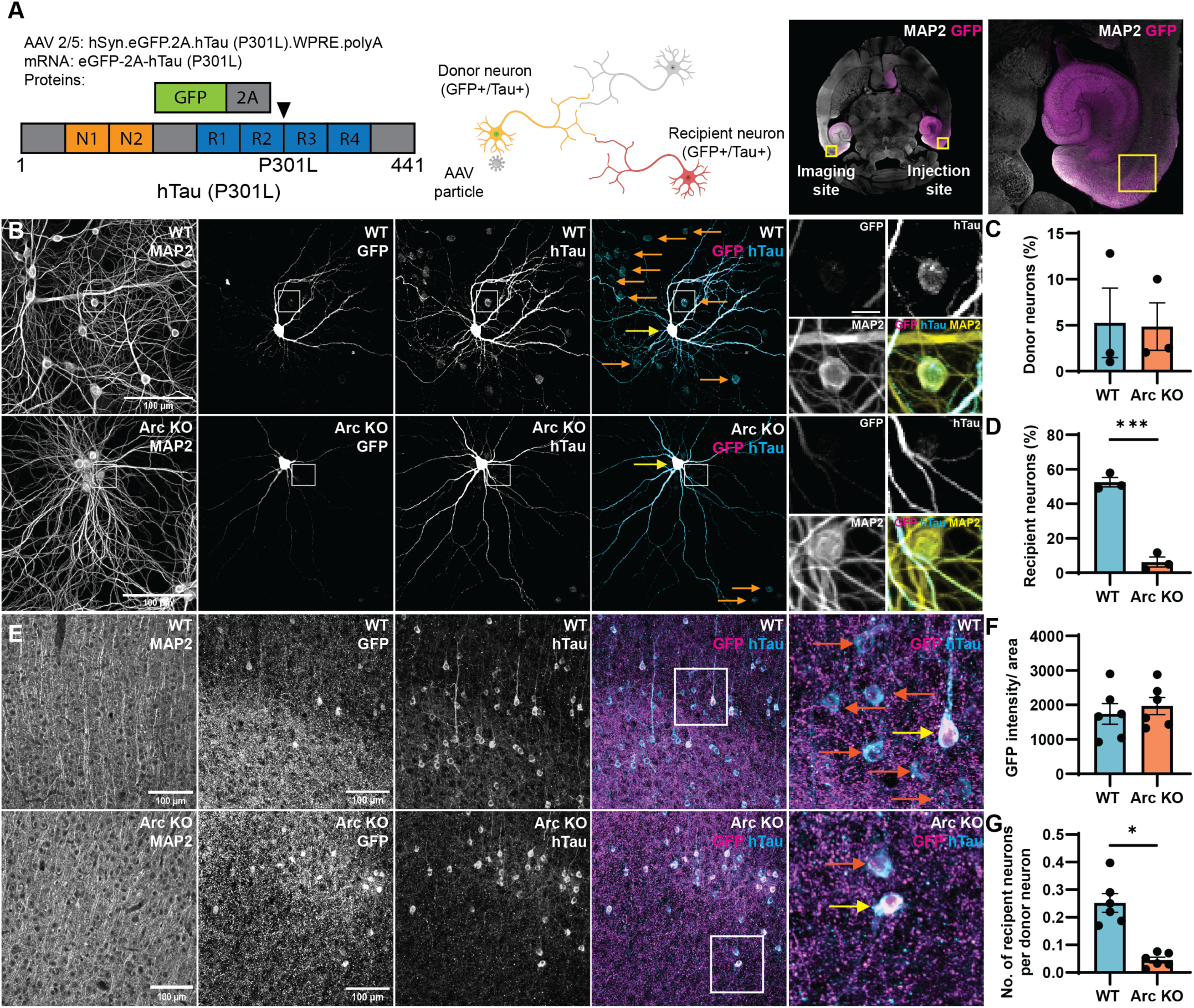
Arc is critical for intercellular hTau transmission. **A.** *Schematic of intercellular hTau transmission assay*. AAV 2/5:hSyn.eGFP.2A.hTau*P301L produces equal amounts of eGFP and hTau (P301L) via a single mRNA cleaved at a P2A sequence. Transduced “donor neurons” express both eGFP and hTau proteins. For *in vivo* studies. 6-month-old WT and Arc KO mice were injected with AAV2/5: hSyn-eGFP-2A-hTau*P301L virus unilaterally in medial entorhinal cortex. Ten weeks post-injection, the brains were collected, sectioned, and immunostained for eGFP and hTau (using Tau Y9 and Tau 13 antibodies). Imaging was performed in contralateral media entorhinal cortex. **B.** *Representative images of hTau and eGFP staining in WT and Arc KO hippocampal primary neurons*. Neurons were sparsely transduced at DIV 7. 14 days post-transduction, the neurons were fixed and immunostained for eGFP and hTau (using Tau Y9 and Tau 13 antibodies). Transduced donor neurons are indicated by yellow arrows and recipient neurons are indicated by orange arrows. **C.** *Virus transduction is the same in WT and Arc KO primary neurons*. The percentage of donor neurons is similar in WT and Arc KO neurons, indicating comparable viral transduction. **D.** *Intercellular hTau transmission is reduced in Arc KO primary neurons.* The percentage of recipient neurons is significantly reduced in Arc KO neurons (n=3 independent cultures). **E***. Representative images of hTau and eGFP staining in WT and Arc KO entorhinal cortex*. Transduced donor neurons are indicated by yellow arrows and recipient neurons are indicated by orange arrows. **F.** *Virus transduction is the same in WT and Arc KO mice entorhinal cortex*. WT and Arc KO mice show no significant differences in eGFP intensity normalized to area, indicating comparable levels of virus transduction in WT and Arc KO mice (n=6, 3M, 3F). **G.** *Intercellular hTau transmission is reduced in Arc KO mice in vivo.* The number of recipient neurons per donor neuron is significantly reduced in Arc KO mice, indicating reduced intercellular hTau transmission (n=6, 3M, 3F). (Statistical analysis: Unpaired t-test, *p<0.05, ***p<0.001).

To test if Arc is important for intercellular hTau transmission *in vivo*, we injected hSyn-eGFP-2A-hTau*P301L in the medial entorhinal cortex of 6-month-old WT and Arc KO mice (Fig. 6A) and collected the brains 10 weeks after injections. We imaged serial sections of the entorhinal cortex contralateral to the injection site where transduction was sparse (Fig. 6A), which allowed us to measure hTau transmission. We imaged and analyzed four 30μm slices from each animal, consisting of 30 images in 1μm steps to obtain a z-stack (see methods). Neurons were classified as eGFP or hTau-positive if values were higher than background values obtained from PBS-injected mice. We observed that eGFP expression, an indication of virus transduction, was comparable between WT and Arc KO mice (Fig. 6F). However, intercellular transmission of hTau was significantly reduced in Arc KO mice *in vivo* (Fig. 6G). Together, these data show that Arc plays a critical role in intercellular transmission of hTau.

**Supplementary Figure 7.**
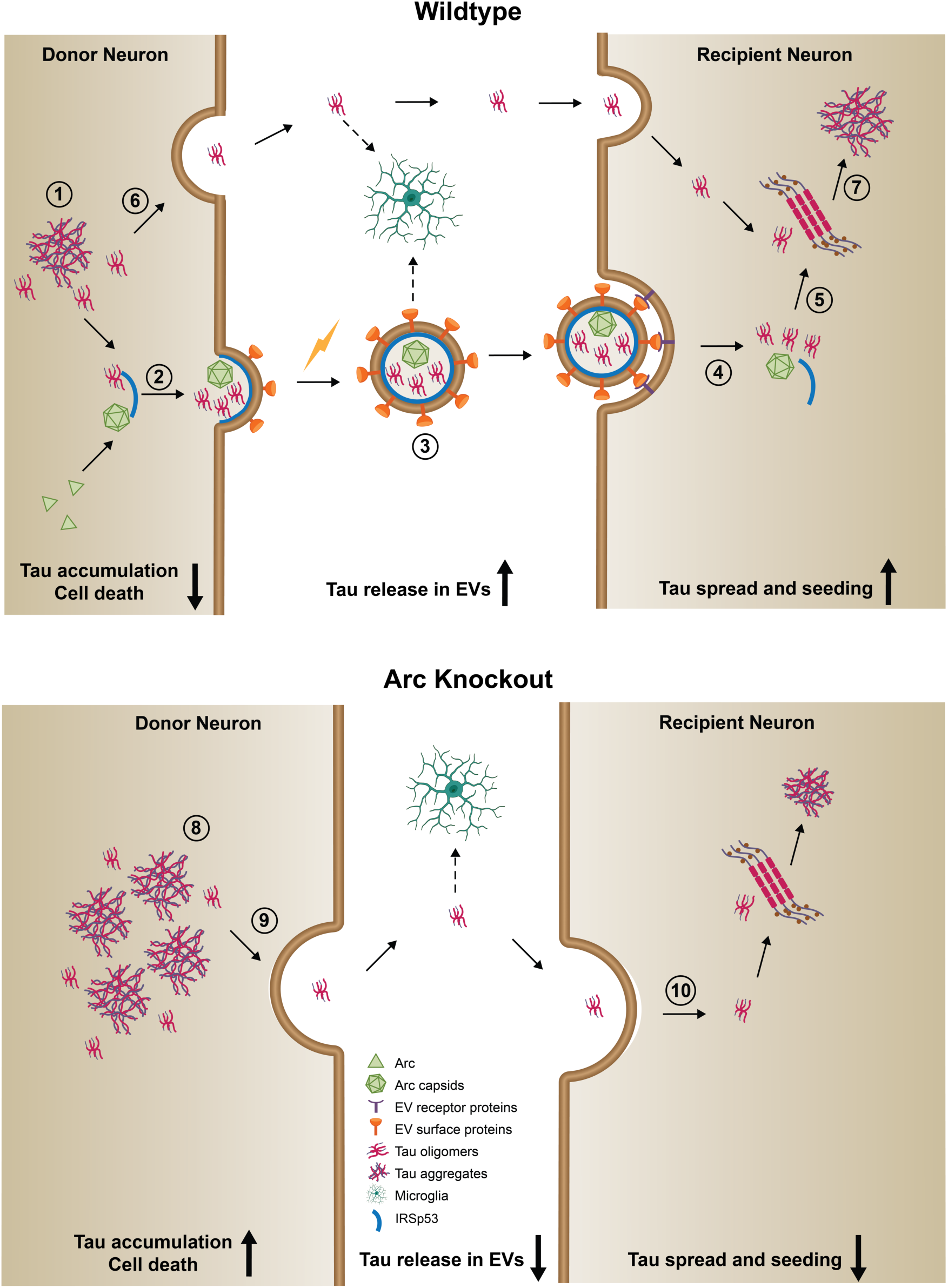
Model of Arc-dependent tau release and intercellular transmission. *Left.* **1.** Early in AD, tau becomes hyperphosphorylated and mislocalized to post-synaptic compartments. **2.** Tau, Arc, and IRSp53 interact, which results in Arc capsid formation and EV biogenesis. **3.** Tau is packaged into Arc-IRSp53 EVs and released in response to neuronal activity. Tau in EVs may be taken up by microglia (dotted arrow). **4.** Arc-IRSp53 EVs are taken up by synaptically connected neurons. **5.** Transmitted misfolded tau seeds intracellular tau aggregation. **6.** Free naked tau is also released from donor neurons. **7.** Free tau is taken up by recipient neurons, or microglia, and contributes to the seeding of intracellular tau aggregation. *Right.* In Arc KO mice: **8.** Misfolded tau accumulates in neurons due to a lack of EV-tau release, which causes cellular toxicity and accelerates cell death. **9.** Free tau is still released and taken up by neurons or microglia. **10.** Overall tau seeding, and spread of pathology, is decreased without intercellular transmission of tau in EVs.

## Discussion

The molecular mechanisms of intercellular tau transmission, despite the importance to understanding AD and FTD pathogenesis, are still unclear. Here, we describe a specific molecular role for Arc in packaging tau into EVs through a direct protein-protein interaction. Arc-mediated release of EV-tau, which we show is capable of tau seeding, plays an important role in both the elimination of intracellular tau and intercellular tau transmission (Fig. S7). These data reveal a detailed biochemical mechanism specific for EV-tau release, independent of free tau release, which requires the biogenesis of Arc EVs via the I-BAR domain protein IRSp53. The profound deficits we observed in intercellular tau transmission in Arc KO mice and neurons suggests that EV-tau is critical for efficient cell-to-cell transmission of pathological tau.

### Packaging of tau in EVs

EV intercellular signaling has emerged as an important component of the spread of pathology in neurodegenerative disorders^64^. However, the function and specific type of neuronal EVs involved in the spread of pathology is poorly described, and their role in neurological disorders is unclear^65^. This is partly due to a lack of mechanistic understanding of the biogenesis and cargo of specific EVs, which are varied and cell-type specific. Endogenous EV-tau release is modulated by neuronal activity and high neuronal activity exacerbates tau pathology^12,21^, but the exact type of EV critical for tau release has remained poorly defined. Arc is released in small EVs (∼100-150nm) that may be released via a classic exosome pathway derived from multi-vesicular bodies^66^. We found that Arc is released from dendrites directly from the cell surface, which requires the I-BAR protein IRSp53^43^. Here we show that hTau is specifically released in Arc-IRSp53 EVs, providing a plausible mechanism for tau packaging into neuronal EVs. Moreover, we also observe Arc and hTau in human brain-derived EVs, suggesting that similar processes occur in the human brain.

Tau, which is normally localized to axons, is mislocalized in dendrites/post synaptic compartments when it becomes hyper-phosphorylated^40,41^. Hyperphosphorylated tau is found at synapses in human AD tissue, both at pre- and post-synaptic compartments^41^. Emerging data also shows that tau is normally locally translated in dendrites^67^ and may be found in the post-synaptic density^51^. We find that hTau forms a complex with Arc and IRSp53 in both mouse and human cortex, consistent with the hypothesis that Arc packages tau into EVs, potentially in dendrites. We found that Arc EV release occurs from dendrites^43^. However, Arc protein has also been identified at presynaptic terminals^68^. Thus, Arc-dependent EV-hTau release may mediate post- to pre-synaptic or post- to post-synaptic tau transmission.

### Tau pathology and intercellular transmission

Our data supports a model where Arc eliminates toxic misfolded intracellular hTau from neurons, which may ultimately prevent cell death. However, a caveat of using hTau transgenic mice is that hTau expression is extremely high. Despite this transgene driven over-expression, we still see clear differences in intracellular hTau accumulation in rTg^Arc^ ^KO^ mice, suggesting a substantial contribution of Arc-dependent elimination of hTau. Our molecular modeling suggests that binding of monomeric hTau to Arc may limit the exposure of aggregation prone elements, which otherwise can seed Tau aggregation^56^, especially in the P301L mutant. The dynamics and fuzzy nature of the Arc-hTau complex may also limit hTau self-assembly inside the cell. In addition to its interactions with monomeric hTau, Arc may also interact with fibrillar hTau through polyelectrolyte interactions with the hTau fuzzy coat ^69^. This may explain our observations that EV-hTau is still capable of seeding aggregation and is consistent with earlier reports that intercellular tau transmission does not depend on its aggregation propensity^70^.

Arc has been implicated in the generation of β-amyloid peptide, by regulating the endocytosis of APP and PS1 in dendrites^31^. APP/PS1 transgenic mice crossed with Arc KO mice show a reduction in amyloid plaques^31^. Thus, Arc may play dual roles in amyloid and tau pathology. The former via an intracellular endocytosis pathway and the latter via an intercellular EV pathway. Data in human AD patients and AD mouse models suggest that neuronal circuits are hyperexcitable early in AD pathogenesis^32,35^, but become hypoexcitable as tau related pathological changes accumulate^71^. In a *Drosophila* tauopathy model, dArc1 expression is increased and reducing dArc1 levels led to decreased neurodegeneration^39^. How dArc1 modulates tau-induced neurodegeneration is unclear and it remains to be determined whether dArc1 intercellular signaling or EV biogenesis regulates tau pathology. Intriguingly, recent studies have implicated endogenous retroviruses (ERVs) in the spread of TDP43 pathology in flies^72^ and tau pathology in cells^73^, which we speculate may also occur through ERV Gag capsid interactions and virus-like particles.

While Arc-dependent release of hTau in EVs may be beneficial, by eliminating intracellular toxic hTau, intercellular hTau transmission is dependent on the balance of release and uptake mechanisms^61^. EV-hTau may first be taken up by microglia, as a mechanism to ultimately degrade toxic hTau^16^. We do not find any overt differences in microglia numbers or activation in the absence of Arc, but this does not preclude a role of microglia in EV-tau transmission. We observed substantial deficits in intercellular hTau transmission in Arc KO neurons *in vitro* and in Arc KO mice *in vivo*. In addition, EVs derived from rTg^Arc^ ^KO^ exhibit decreased hTau seeding potential, which suggests that Arc packaging and release of EV-hTau plays a critical role in intercellular tau transmission and possibly the spread of tau pathology. The mechanisms that are most important for the uptake of monomeric tau, such as through LRP receptors, are likely not as efficient for phosphorylated tau^74^ and EV-tau uptake may be more efficient than naked tau^4^. Thus, EV-tau uptake may play an important role in neuron-to-neuron transmission of bioactive tau as we demonstrate here. Our findings suggest that anti-tau antibody therapy may not prevent all forms of tau transmission as EVs may preclude antibody binding. Understanding the mechanisms of EV-tau transmission may provide novel targets for preventing the spread of tau pathology.

## Conclusion

Together, our data suggests a model (Fig. S7) where misfolded tau binds Arc to facilitate packaging of tau in EVs. EV-tau is either taken up by neurons or microglia. While Arc-dependent EV-tau release may help prevent cell death due to toxic accumulation of misfolded tau, excessive EV-tau release, perhaps due to hyperexcitable neurons, contributes to the intercellular transmission of tau and eventually the spread of tau pathology.

## Acknowledgements

This work was supported by a NIH Director’s Office Transformative Research Award (R01 NS115716), the Chan-Zuckerberg Initiative Ben Barres Early Acceleration Award, the McKnight Brain Disorders Award, and the Jon M. Huntsman Presidential Endowed Chair fund (J.D.S); NINDS DSPAN F99 (K.S). AIRC IG 26229 and PRIN 2022EMZJL4 (M.F.). The Rainwater Foundation, the JPB Foundation, Cure Alzheimer Fund, and NIH grant AG073236 (B.H). The Massachusetts Alzheimer’s Disease Research Center (P30AG062421) provided human samples.

We would like to thank Kaylyn Bauer in Dr. Ryan O’Connell’s lab for assisting with NTA experiments. We thank Dr. Kenneth Lyon for generating custom Mathematica code for image analysis. We thank the University of Utah Flow Cytometry and imaging core facility and the University of Utah Drug Discovery Core for AAV generation. We thank Alyson Stewart for generating primary cultured neurons and Alex Holbrook for animal colony management. We also thank Karla McHale and the Shepherd lab for supporting these studies.

## Author Contributions

M.T. and J.D.S. conceived the project. M.T. and J.D.S. wrote the manuscript; all authors discussed results and edited the manuscript. M.T. performed and analyzed tau transfer assays, immunoprecipitation experiments, proteinase K protection assays, tau seeding assays, ELISA assays, GST pull-down assays, western blots, and immunohistochemistry of rTg mice. R.C. developed the mouse brain EV isolation protocol, conducted NTA assays, performed western blot analyses to characterize EVs and compare rTg^WT^ and rTg^Arc^ ^KO^ mouse EVs, generated 3XTg^Arc^ ^KO^ mice, and conducted immunohistochemistry of 3XTg AD mice. E.D. performed and analyzed electrophysiological experiments and the hTau HiBit assay. K.S. performed protein purification, electron microscopy, and plasmid cloning. A.W. sliced mice brains and helped conduct brain EV isolation from mouse tissue. A.N. helped perform and analyze microglia immunohistochemistry data. B.F. and M.F. performed computational modeling, and M.F conceptualized the Arc-hTau interaction. B.H. provided reagents and human brain tissue samples. J.D obtained funding for the project.

## Methods

### Plasmids

hSyn-eGFP-2A-human WT hTau and hSyn-eGFP-2A-human P301L hTau plasmids were previously generated in the Hyman lab^63^. Human P301L tau was amplified by PCR and ligated into the C-terminal pLVX-HiBiT vector to generate pLVX-human tau-HiBiT. The open reading frame (ORF) of full-length rat Arc (NP_062234.1) was amplified by PCR and ligated into the pRK5 vector to generate pRK5-Arc. Similarly, the ORF of mouse IRSp53 (transcript variant 1, NM_001037755) was inserted into the pLVX lentiviral vector. For protein purification, the pGEX-6p1 bacterial expression vector (GE Healthcare) was used. The ORFs of human tau (WT and P301L, rat Arc, and mouse endophilin were amplified by PCR and subsequently ligated into the pGEX-6p1 GST vector.

### Animals

All procedures were performed following the guidelines of the Institutional Animal Care and Use Committee of the University of Utah. All mice were housed in breeding pairs, or group-housed with littermates of the same sex after weaning (2-5 mice/cage), on a 12:12 h day:night cycle, with food and water provided *ad libitum*. Both male and females were used for all studies.

C57BL/6 Arc germline knock-out (KO) mice^36^ were crossed with rTg4510 hTau transgenic mice obtained from the Jackson Laboratory (Strain: 015815). The animals were backcrossed for > 5 generations. Mice heterozygous for hTau transgenes and Arc genes were crossed to generate rTg^WT^ and rTg^Arc^ ^KO^ littermates. We obtained heterozygous 3xTg AD mice from Jackson Laboratory (strain: 034830). F1 offspring were genotyped to confirm heterozygosity and were bred together to generate homozygous 3xTg mice. Homozygous 3xTg mice were then crossed with Arc KO mice to obtain 3xTg^Arc^ ^het^ mice. 3xTg^Arc^ ^het^ mice were interbred to generate 3xTg^Arc^ ^WT^ and 3xTg^Arc^ ^KO^ mice.

### Antibodies

Western blot – Primary antibodies: Tau 13 (1:1000, anti-mouse, Biolegend), HT7 (1:1000, anti-mouse, Thermo Fisher), TauY9 (1:1000, anti-rabbit, Enzo Life Sciences), T22 (1:1000, anti-rabbit, EMD Millipore), AT8 (1:1000, anti-mouse, Thermo Fisher), Arc (1:1000, anti-rabbit, Synaptic systems), IRSp53 (1:1000, anti-mouse, Abcepta), Syntenin (1:500, anti-rabbit, Abcam), Alix (1:500; rabbit polyclonal, custom provided by the Dr. Wesley Sundquist), CD9 (1:500, anti-rabbit, Abcam), Histone 3 (1:500, anti-rabbit, Abcam). Secondary antibodies: All HRP conjugated secondary antibodies (Jackson ImmunoReseacrh) were used at 1:5000 dilution.

*Immunostaining: Primary antibodies:* GFP (1:1000, anti-chicken, Abcam), Tau 13 (1:1000, anti-mouse, Biolegend), TauY9 (1:1000, anti-rabbit, Enzo Life Sciences), NeuN (1:500, anti-guinea pig, EMD Millipore), MAP2 (1:500, anti-chicken, abcam), and Iba1 (1:1000, anti-rabbit, Wako). Secondary antibodies: Alexa Fluor 405, 488, 555, or 647 (1:1000 Thermofisher Scientific) secondary antibodies.

### Intracranial virus injections

Animals were deeply anesthetized with isoflurane (3% for induction, 1.5–2% for surgery) and immobilized in a mouse stereotaxic frame (David Kopf Instruments) on 37°C warming pad. Before placing the ear bars, the antibiotic Baytril (8 mg/kg, VetOne), the analgesic Carprofen (5 mg/kg, Zoetis) and the steroid dexamethasone (13 mg/kg, VetOne) were administered. Mice were head-fixed using the ear bars and Lidocaine (100 mg/ kg; VetOne) was injected subcutaneously beneath the scalp. A midline incision was made to expose the skull, and a small burr hole was made with a pneumatic dental drill above the EC (relative to bregma: anterior/posterior (A/P): −4.7 mm, medial/lateral (M/L): ±3.3 mm). A glass pipette filled with virus, was lowered (−2.0 mm from brain surface) and allowed to rest for 5 min. Unilateral injection of AAV2/5:hSyn-eGFP-2A-huTau (P301L) (2.9×10^9^ particles, generated by the University of Utah Drug Discovery Core) was delivered using a Nanoject II AutoNanoliter Injector (Drummond Scientific). After an additional 5 min of rest following the injection to allow diffusion of virus, the pipette was slowly removed. The scalp was closed using sutures (Ethicon) and the animals were allowed to recover from anesthesia on 37°C warming pad. The mice were returned to clean cages for recovery and were monitored for signs of infection or discomfort for 3 days.

### Primary neuron cultures

Primary hippocampal and cortical neuron cultures were prepared from E18 mouse embryos as previously described^53^. Cortices and hippocampi from embryos were dissociated using 0.067% papain (Worthington Biochemicals) and 0.01% DNase (Sigma-Aldrich), followed by serial trituration with fire-polished glass pipettes to obtain single cell-suspension. Cells were pelleted by centrifuging at 500g for 4 mins, the supernatant was discarded, and the cells were resuspended in neurobasal media (Thermo Fisher) containing 5% fetal bovine serum (FBS; Thermo Fisher), 1% glutamax (Thermo Fisher), 1% penicillin/streptomycin (P/S; Thermo Fisher), and 2% SM1 (StemCell Technologies). The cells were then plated on coverslips (Carolina) in 12-well plate at 1.8×10^5^ cells/well or in 10 cm culture dishes at 6×10^6^ cells/dish, both coated with 0.2mg/ml poly-L-lysine (Sigma-Aldrich) overnight. Neurons were cultured at 37°C with 5% CO_2_ and fed every third day by half-media exchange using astrocyte-conditioned BrainPhys media (StemCell Technologies) supplemented with 1% FBS, 500 µM L-Glutamine (Gibco), 1% P/S, and SM1. On *days in vitro* (DIV) 4, 5µM Cytosine β-D-arabinofuranoside (Sigma-Aldrich) was added to inhibit glial cell proliferation and obtain relatively pure neuron cultures.

### Mouse brain homogenate preparation

Forebrains was dissected and washed with 1x HBSS. The brains were homogenized in ice-cold 2ml 0.5% IP buffer (150 mM NaCl, 50 mM Tris, 0.5% Triton X-100, pH 7.4) containing protease inhibitor (Roche) and phosphatase inhibitor (Roche) using a Dounce homogenizer. Homogenates were centrifuged at 200g for 5 min at 4 °C, followed by centrifugation at 13,000g for 10 min at 4°C. The final supernatant was collected, adjusted to a volume of 2 ml, and used for western blot analysis

### Western blot

Protein samples for western blot analysis were mixed with 4X Laemmli buffer (40% glycerol, 250 mM Tris, 4% SDS, 50 mM DTT, pH 6.8), heated at 95°C for 5 mins, and separated on a 10% SDS-PAGE gel. The gels were transferred to a nitrocellulose membrane (GE Healthcare) through a wet transfer at 100V for 1 hr. Membranes were stained with Pierce™ Reversible Total Protein Stain and then destained. Membranes were blocked with 5% milk in 1x TBS for 1 hr at RT. Primary antibodies were diluted in 1% milk in 1x TBS and incubated with the membrane at 4°C overnight. Membranes were washed 3 x 10 min in 1X TBST (0.1% Tween-20 in 1x TBS) and then incubated with HRP-conjugated secondary antibodies (Jackson ImmunoResearch) diluted in 1% milk in 1x TBS for 1 hr at RT. Membranes were then washed 3 x 10 min in 1X TBST. Protein bands were visualized using Clarity™ Western ECL Substrate (Bio-Rad) and imaged using an Amersham ImageQuant™ 800 imaging system (Cytiva). Images were analyzed and quantified using ImageJ (National Institutes of Health).

### Protein purification

GST-tagged proteins were purified from *Escherichia coli* Rosetta 2 BL21 competent cells as previously described^36^. Starter bacteria cultures were grown overnight at 37°C in LB supplemented with ampicillin and chloramphenicol. Large-scale 500 mL cultures were inoculated with starter cultures in ZY auto-induction media. Large-scale cultures were grown to OD_600_ of 0.6-0.8 at 37°C at 160rpm then shifted to 18°C at 180rpm for 16-20 hours. Cultures were pelleted at 4000*g* for 15 min at 4°C. Pellets were resuspended in 25 mL of lysis buffer (500 mM NaCl, 50 mM Tris, 5% glycerol, 1 mM DTT, pH 8.0) and flash frozen in liquid nitrogen. Frozen pellets were thawed in 37°C and brought to a final volume of 1g pellet: 10ml lysis buffer, supplemented with DNAse, lysozyme, and complete protease inhibitor cocktail (Roche). Resuspended lysates were sonicated for 6 x 45s pulses at 90% duty cycle and pelleted for 75 min at 21,000*g*. Cleared supernatants were incubated with pre-equilibrated GST Sepharose 4B affinity resin in a gravity flow column overnight at 4°C. GST-bound protein was washed twice with 20 resin bed volumes of lysis buffer and re-equilibrated in TBS (150 mM NaCl, 50 mM Tris, 1 mM EDTA, 1mM DTT, pH 7.2) at RT, and cleaved from GST resin overnight at 4°C with PreScission Protease (GE Healthcare). GST was affinity purified as described above and eluted using 15 mM reduced L-gluathione, 10 mM Tris, pH 7.4 at RT.

### GST pull-down assays – HEK293 cells

9µg of hSyn-eGFP-2A-human WT or P301L tau plasmid was expressed in 10 cm biochemical dish of HEK293 cells for 24 hours. 1ml of 0.5% IP buffer containing protease inhibitor and phosphatase inhibitor was added and cell lysates were collected. The cell lysates were centrifuged at 200g for 10 min. and supernatant was collected. The GST-tagged proteins immobilized on beads were washed 3 times with wash buffer (50mM Tris, 150mM NaCl, 1mM EDTA, pH 7.2). 0.2 mg of GST-tagged proteins immobilized on beads were incubated with equal amounts of untransfected, WT tau transfected, and P301L tau transfected HEK cell lysates overnight at 4°C. The beads were loaded on gravity flow columns (Biorad) and washed 3 times with 10 ml wash buffer. GST-tagged proteins were eluted from Sepharose beads by incubating with 500ul of cleavage buffer (20mM L-reduced glutathione, 50mM Tris, pH 8.0) for 20 minutes at room temperature. The elutes were collected and blotted for tau using HT7 and Tau 13 antibodies.

### GST pull-down assays – purified proteins

0.2 mg of GST or GST-Arc was purified and immobilized on beads. 0.2 mg of recombinant full-length human mutant (P301L) tau (Abcam) was incubated with immobilized GST or GST-Arc overnight at 4°C. The beads were loaded on gravity flow columns (Biorad) and washed 3 times with 10 ml wash buffer. GST-tagged proteins were eluted from Sepharose beads by incubating with 500ul of cleavage buffer (20mM L-reduced glutathione, 50mM Tris, pH 8.0) for 20 minutes at room temperature. The elutes were collected and blotted for tau using HT7 and Tau 13 antibodies.

### Immunoprecipitation

Mouse forebrain was dissected and homogenized in 2ml of IP lysis buffer (150 mM NaCl, 50 mM Tris, 0.3% Triton X-100, pH 7.4) with freshly added protease inhibitor cocktail (Roche) and phosphatase inhibitor (Roche). The homogenates were centrifuged at 200g for 5 min at 4°C to remove tissue debris. The supernatant was collected, mixed, and then centrifuged at 10,000g for 10 min at 4°C to remove insoluble material. The clarified supernatant was collected, and divided into input (10%), IgG (45%), and IP (45%) samples. Supernatants were immunoprecipitated with either Arc antibody (rabbit polyclonal, synaptic systems) or normal rabbit IgG (rabbit polyclonal, EMD Millipore) at 2 µg/ml overnight at 4°C with rotation. Protein A/G magnetic beads (Pierce) were washed 3 times with IP buffer and added to the antibody-supernatant mixture (10% of the total volume). The mixture was rotated for 1 hr at 4°C. The beads were pulled out using a magnetic stand, washed three times with IP buffer, and finally resuspended in 200 µL IP buffer. To cleave proteins from magnetic beads, 4X Laemmli buffer was added, and samples were heated 70°C for 5 mins. Proteins were separated from magnetic beads using a magnetic stand. For detection, Tau 13 (1:1000, anti-mouse, Biolegend), Arc E5 nanobody^75^ (40ug, anti-alpaca, purified in house and IRSp53 (1:1000, anti-mouse, Abcepta) were used. For human IPs, 0.1g of frozen post-mortem tissue was homogenized in 1ml of 0.3% IP buffer. The homogenates were processed as above. Supernatants were immunoprecipitated with either D5D8N hTau antibody (rabbit mouse antibody, Cell Signaling) or D1M9X normal IgG (rabbit mouse antibody, Cell Signaling) at 2 µg/ml overnight at 4°C with rotation. For detection, Tau 13 (1:1000, anti-mouse, Biolegend), Arc (1:200, anti-mouse, Santa Cruz), and IRSp53 (1:1000, anti-mouse, Abcepta) were used.

### Negative-stain electron microscopy

Copper 200-mesh formvar/carbon coated grids (Electron Microscopy Sciences) were discharged for 25 seconds in a vacuum chamber. EVs were fixed at room temperature for 30 minutes using 0.1% paraformaldehyde in PBS. EVs were added at 0.3mg/ml to the discharged grids and incubated for 45 seconds at RT^76^. Grids were then washed 3 times with 30µl distilled water for 5 seconds. The grids were then washed once with 20µl 1% uranyl acetate for 5 seconds and stained for 30 seconds. The excess solution was removed using filter paper, and the grids were air-dried for 1 minute. The grids were imaged using a JEOL-JEM Transmission Electron Microscope operated at 120kV. 15 Images were taken at predetermined grid locations at 20,000X for all samples.

### Isolation of extracellular vesicles from mouse brain tissue

Whole mouse brains, excluding the cerebellum, were flash-frozen immediately after extraction. Brain tissue from 2 mice were used for each EV preparation. Brain tissue was rinsed with 0.2µm filtered phosphate buffer saline (PBS) and sliced into 2mm^3^ cubes using a sterile scalpel blade in modified RPMI-1640 (Sigma) on ice. Tissue pieces were then moved to a 6-well plate and incubated with Collagenase D (0.24U/mg, Roche) and DNase I (10U/µl, Sigma) for 30 minutes at 37°C and shaken at 60 rpm. The resulting suspension was filtered through a 70µm cell strainer (Corning) and collected in a 50ml falcon. Filtered media was sequentially spun at 300g for 10 minutes and 2000g for 20 minutes at 4°C. The supernatant was transferred to a round bottom tube using a 20ml plastic syringe with a blunt needle (2.10mm x 80mm) and spun at 16,500g for 20 minutes at 4°C (45 Ti fixed angle titanium rotor) to pellet larger vesicles and apoptotic bodies. The clarified supernatant was transferred to another tube and spun at 24,000g for 16 hours at 4°C to obtain small extracellular vesicles. This pellet was dissolved in 300µl PBS and further fractionated using qEV 35nm IZON size exclusion columns (IZON). Eleven 0.5ml fractions were collected and early fractions (1-4) were combined and concentrated to 120µl using 100kDa Vivaspin protein concentrator spin columns (Cytiva) to obtain final EV samples.

### Proteinase K protection assay

Brain EVs were isolated from three 4-month-old rTg^WT^ mice as above. Three conditions were set up (total volume of 27ul): 1. 20ug of EVs + 7ul water 2. 20ug of EVs + 6µL water + 1µL proteinase K (200µg/mL) (New England BioLabs) 3. 20ug of EVs + 3µL water + 3µL 10% Triton X-100 + 1µL proteinase K (200µg/mL). The EVs were incubated for 10 min. at RT and then 3 µL of PMSF (10 mM) was added. Samples were mixed with 4x Laemmli buffer, boiled at 95°C for 5 min., and used for western blot analysis.

### Nanoparticle tracking analysis

Size and concentration of EV fractions were determined using a Nanosight NS300 (Malvern Technologies). 5µl of each sample was diluted in 700µl PBS. For each sample, three 60s videos were taken at 25 frames per second. The same detection threshold was maintained for each sample to ensure consistent measurement of particles.

### HEK biosensor cell tau seeding assay

HEK biosensor cells stably expressing P301S hTau repeat domain conjugated to either cyan fluorescent protein (CFP) or yellow fluorescent protein (YFP) (hTau^RD^-P301SCFP/YFP) were obtained from ATCC (CRL-3275). Cells were maintained at 37°C/5% CO2 in DMEM media (Thermo Fisher) supplemented with 10% FBS and 1% penicillin-streptomycin (Thermo Fisher). Poly-D-lysine coated 96-well cell culture treated plate (Costar) was plated with 90,000 cells per well and cultured for 24 hours. 10µg of EVs diluted in opti-MEM (Thermo Fisher) were mixed with 1.25 µl lipofectamine 2000 (Thermo Fisher) for a total volume of 20ul and incubated for 20 min at RT. The cells were incubated with the EV-lipofectamine mixture for 24 hours. The cells were then harvested with 0.05% trypsin (Gibco). Trypsin was inactivated using DMEM and cells were fixed in 2% paraformaldehyde (Thermo Fisher) for 10 min. The cells were resuspended in 1x PBS and used for flow cytometry analysis.

### FRET flow cytometry

A Beckman Dickinson FACS Canto II (BD Biosciences) was used to perform FRET flow cytometry. The cells were excited 405 nm and 488 nm lasers for CFP/FRET and YFP, respectively. CFP and FRET fluorescence was captured using 405/50 nm and 525/50 nm filter, respectively. YFP fluorescence was captured using 525/50 nm filter. A bivariate plot of FRET versus CFP was created, and a triangular gating strategy was applied to identify and quantify the population of cells that exhibited FRET-positive signals. The FRET gate was calibrated based on biosensor cells treated with lipofectamine only, which are expected to be FRET-negative, ensuring accurate distinction between positive and negative FRET signal. FACSDiva software vs 6.3 (BD Biosciences) was used for analysis to calculate the number of FRET positive cells and median fluorescence of FRET-positive cells. Integrated FRET density, calculated as the percentage of FRET-positive cells multiplied by their mean fluorescence intensity was calculated.

### Neuron culture tau release assay

WT and Arc KO cultures were transduced with equal amounts of AAV2/5: hSyn-eGFP-2A-P301L tau on DIV 7. A full media change with BP1+SM1 was performed on DIV 16. Conditioned media was collected on DIV 18. The media was centrifuged at 200g for 5 min at 4°C to remove cell debris. The supernatant was collected and centrifuged at 2000g for 10 min at 4°C. The supernatant was concentrated 70 times using 30KDa vivaspin 20 concentration column (Cytiva). The concentrated media was divided into two parts and mixed with PBS or 1% Triton-X-100 detergent. EVs were incubated with PBS or detergent for 10 min at RT. The hTau in PBS or detergent treated media was quantified using total human tau ELISA (Invitrogen).

### Tau Hibit assay

Neuro-2A (N2A) cells were obtained from ATCC (CRL-11268). Cells were maintained at 37°C/5% CO2 in DMEM media supplemented with 10% fetal bovine serum (FBS) (Thermo Fisher). N2A cells were transfected at 70% confluency in 6-well plates. Cells were transfected using Polyethylenimine (PEI) (1mg/mL) (Polysciences). 3µg of plasmid DNA was diluted in OptiMEM. DNA and PEI were incubated for 20 min at RT and incubated with cells overnight at 37°C. A full media change was then performed with DMEM supplemented without FBS and conditioned media was collected 24 hours later. For hTau-HiBiT experiments, N2A cells were transfected with pLVX-TauP301L-HiBiT with pRK5-Arc, and/or pLVX-IRSp53. Conditioned media was centrifuged at 1000g for 10 min and supernatant was either suspended in Passive Lysis buffer (Promega) (total hTau) or the equivalent amount of PBS (free hTau). Media containing Passive Lysis buffer was then freeze-thawed to break open EVs. Tau-HiBiT signal was measured by complementation with LgBiT (equal amounts and measurement of Nanoluc-mediated luciferase luminescence (Nano-Glo Luciferase Assay, Promega, 565nm) on a microplate reader (SpectraMax iD3). Total tau in the cell lysate and the conditioned media was determined by dividing the luminescence value by the proportion of sample used to total sample collected. % EV tau release was calculated by subtracting naked tau from total tau and dividing by tau in the cell lysate.

### Electrophysiology

Current clamp electrophysiological recordings of cultured hippocampal neurons: Whole-cell patch clamp recordings were obtained from wildtype and *Arc* knockout DIV 16 hippocampal neurons constantly perfused with aCSF containing (in mM) 125 NaCl, 2.5 KCl, 26 NaHCO_3_, 1.25 NaH_2_PO_4_, 2 MgSO_4_, 2 CaCl_2_, 10 Glucose, 10 HEPES (pH 7.4; 295 mOsm) at a temperature of 31-33 °C with a flow rate of 1.5 ml/min. Gigaohm seal formation was achieved with patch pipettes (4-6 MΩ) filled with internal solution containing (in mM) 120 K-gluconate, 20 KCl, 2 MgCl_2_, 10 HEPES, 7 phosphocreatine, 0.2 EGTA, 4 Na_2_-ATP, 0.3 Na_2_-GTP (pH 7.2; 280-290 mOsm). Following seal formation, patch pipette capacitance compensation, whole cell breakthrough, and pipette resistance compensation (with a bridge balance circuit) was achieved. Access resistance was maintained at σ; 30 MΩ. Recordings were obtained with a Multiclamp 700B and a Digidata 1440A (Molecular Devices), sampled at 25 kHz, and low-pass filtered at 10kHz. Data analysis was performed with Easy Electrophysiology 2.6.1, Clampfit 11 (Molecular Devices) and Graph Pad Prism 10.

Spontaneous Firing and Excitability: To determine the spontaneous firing activity of cultured hippocampal neurons, we recorded firing activity for 3 minutes and quantified the firing frequency across 3 minutes of spontaneous activity. To assess excitability, neurons were stimulated with a current-step protocol that consisted of 2-second current steps between −200 and +500 pA in 25-pA intervals. To quantify excitability, we determined the firing frequency at current steps between 0 and 500 pA. For spontaneous firing and excitability, statistical significance was determined using an unpaired t-test and two-way ANOVA, respectively.

### Immunohistochemistry

For immunofluorescence labeling of brain sections, mice were transcardially perfused with ice-cold phosphate buffer saline (PBS) followed by PBS containing 4% paraformaldehyde (PFA) (Electron Microscopy Sciences). Brains were extracted, postfixed in 4% PFA/PBS for 1 day at 4°C, and then cryoprotected in 30% sucrose in PBS for 3 days. The brains were horizontally sectioned into 40-μm-thick slices using a cryostat. The floating sections were washed 3×10 min with 1x PBS and blocked with 5% normal donkey serum (Jackson ImmunoResearch), 2% fish gelatin (Thermo Fisher), 0.1% Triton X-100 (Amresco), and 0.05% sodium azide (Sigma-Aldrich) in 1X PBS for 2 hours at RT. Sections were incubated with primary antibodies diluted in the blocking solution overnight at RT. Sections were then washed three times over 24 hours with wash buffer (2% fish gelatin, 0.1% Triton X-100, and 0.05% sodium azide in 1X PBS). Secondary antibodies, diluted in the blocking buffer, were incubated with sections overnight at RT. The sections were washed again three times over 24 hours in wash buffer. After washing, sections were mounted on slides, and coverslips were applied using. The slides were dried flat overnight at room temperature and then stored at 4°C in slide boxes.

### Immunocytochemistry

Neurons were briefly washed once in 1x phosphate-buffered saline (PBS) and then fixed in 4% paraformaldehyde (PFA/4% sucrose in 1x PBS for 15 min at RT. Following fixation, the neurons were washed 3X5 min with PBS, permeabilized with 0.2% Triton X-100 in PBS for 10 min, and washed 3X5min with PBS. The neurons were blocked in 10 mg/mL bovine serum albumin (BSA) in PBS for 30 min and incubated with primary antibodies diluted in blocking buffer for 1 hr at RT. After incubation, the neurons were washed 3X5 min in PBS and incubated with secondary antibodies prepared in the blocking buffer for 1 hr at RT. Following 3X5min washes with PBS, neurons on coverslips were mounted on slides using Fluoromount (Thermo Fisher) and stored flat at RT overnight.

### Post-mortem human tissue

Flash-frozen post-mortem tissue from one control (M, 87years), one Braak stage 2 AD patient (M, 90 years), and one Braak stage 6 AD patient (F, 67 years) was obtained from Massachusetts Alzheimer’s Disease Research Center brain bank. Post-mortem tissue from healthy or AD human participants was selected based on several factors, including clinical diagnosis of dementia due to probable AD, postmortem confirmation of AD diagnosis, and determination of Braak stage pathology by total hTau immunostaining and Bielchowsky’s silver stain. Human brains were collected with informed consent of patients or their relatives and approval of local institutional review boards at Massachusetts General Hospital. Prefrontal cortex corresponding to Broadman Area 8/9 from healthy and AD patients was dissected and stored at −80°C until use. Approval from the IRB at the University of Utah was obtained to carry out human work.

### Isolation of extracellular vesicles from human brain tissue

Human tissue derived EVs were isolated as previously described^4^. 0.5 g of flash-frozen human prefrontal cortical post-mortem tissue was sliced with a razor blade into 2–3 mm^3^ sections in Hibernate-E media (Thermo Fisher) on ice. The tissue sections were dissociated in 3ml Hibernate^TM^-E medium (containing 20 units of papain at 37°C for 15 min). Post-incubation, 6 ml of ice-cold Hibernate^TM^-E media was immediately added. The media was filtered through 40 μm mesh filter (Corning) and centrifuged at 300g for 10 min at 4°C. The supernatant was collected and centrifuged at 2000g for 10 min at 4°C. The supernatant was transferred to round-bottom tubes and centrifuged at 10,000g for 10 min at 4°C. The supernatant was passed through a 0.22 μm filter before undergoing ultracentrifugation at 100,000g for 70 min at 4°C (Optima-XE SW41 Beckman Coulter). The resulting pellet was then resuspended in 2 ml of 0.475 M sucrose solution prepared in double-filtered PBS. This mixture was carefully layered over a sucrose gradient (2 ml each of 2.0 M, 1.5 M, 1 M, 0.825 M, and 0.65 M in dfPBS) and ultracentrifuged again at 200,000g for 20 hrs. at 4°C. The gradient was collected in 2 ml fractions. EV-enriched fractions V and VI were collected, diluted in 12 ml PBS, and ultracentrifuged at 100,000g for 70 min at 4°C. The final pellet was resuspended in 60ul of PBS and used for western blot analysis.

### Computational modelling

Using the sequence of the human Tau (2N4R, Uniprot: P10636-8) four repeat domains (R_1_-R_4_, residues 244-369), we generated models for its complex with mouse (Uniprot: Q9WV31) and human Arc (Uniprot: Q7LC44) proteins using the AlphaFold program^77^. We predicted the structures for both the wild-type Tau sequence, as well as the disease-associated P301L Tau mutant and used three representative wild-type and five mutant Tau conformations as starting models for the simulation. The models were placed into rhombic dodecahedron simulation cell (20 nm edges), were solvated with TIP3Pε waters and ions were added to model 150 mM NaCl. The simulation system contained approximately ∼1 million particles. Using CHARMM36m force field^78^ the system was optimized and heated to 310K while gradually removing positional restraints. The system was equilibrated using molecular dynamics simulations with GROMACS^79^, while monitoring the hydrodynamic radius (Rg). When Rg converged, due to the collapse of the disordered Arc tails, the system was re-solvated. We performed an NPT molecular dynamics simulations using the re-solvated systems with ∼500 000 atoms, with velocity-rescaling thermostat and a Parrinello-Rahman barostat with 2 fs integration steps. For each model a 200 ns simulation was carried out, and the first 100 ns trajectory was discarded. The trajectories of the different models were combined for analysis, which was performed by the ConAn program^80^. The secondary structure populations, in particular the β propensities, computed as the fraction of the combined 500 ns equilibrated trajectory using the STRIDE algorithm^81^. The contacts were defined with 5.5 Å threshold between the closest heavy atoms.

### Imaging and analysis

Imaging was performed using a 20x 0.45 NA objective on a Nikon Eclipse Ti2-E inverted confocal microscope and images were analyzed using ImageJ and Mathematica. The threshold for each experimental group was determined using no primary controls, with image acquisition settings determined by the brightest immunofluorescent sample Tau pathology in rTg^WT^ and rTg^ArcKO^ mice: CA1 region of the dorsal hippocampus was imaged in 2 slices (2 images) for each animal. Imaging was performed through 40µm slices with 1µm step size. The number of NeuN-positive neurons and the average tau expression per neuron was quantified using our custom Mathematica analysis pipeline. We employed our analysis pipeline to generate a NeuN mask and use local binarization to detect all neurons in each z-plane of the z-stack. The pipeline defined a region of interest (ROI) for each detected neuron. The ROIs were manually checked and corrected for any errors. Since each neuron is counted 6-8 times in a 3D volume due to average size of a neuron being 6-8μm^82^, the overall number of NeuN positive neurons were divided by 7 to obtain an average value for the z-stack volume. The ROIs were overlaid with the hTau channel, and the hTau raw integrated density/area for each NeuN-positive neuron was calculated. A cutoff for hTau-positive neurons was established using the no primary control. The hTau raw integrated density/area was averaged for all hTau-positive neurons. The value obtained from each image were averaged to derive a single representative value for each animal.

Tau transfer assay in cultured neurons: 5µm images were acquired with a step size of 0.25 µm. Fields of view around eGFP-positive neurons were imaged. ROIs were created using Image J for all neurons within the MAP2 channel, agnostic to eGFP and hTau staining. The ROIs were transferred to eGFP and hTau channels and the raw integrated density/area values were calculated for each channel. Neurons were classified as eGFP-positive and hTau-positive if their raw integrated density/area values were higher than background values of neurons in untransduced controls. Percentage of donor or recipient neurons were calculated for each image and averaged across 5 images to get an experimental value. Values for three cultures were averaged to get the final plot.

Tau transfer assay in vivo: Medial entorhinal cortex contralateral to the injection site was imaged and analyzed. Imaging was performed through four 30 µm slices per animal, including two dorsal and two ventral hippocampal slices, with a 1 µm step size in each slice. An overlay of GFP and tau z-stack images was made to generate a mask. Our custom analysis pipeline was then used to detect all neurons using local binarization and define ROIs for all detected neurons. The ROIs were transferred to GFP and hTau channel and raw integrated density/area values were calculated for each channel. A cutoff for classifying eGFP-positive and hTau-positive neurons was established using images from PBS-injected mice. The number of donor and recipient neurons was calculated by adding the counts from each z-plane. The number of donor and recipient neurons were divided by 7. The values obtained from each image were averaged to derive a single representative value for each animal.

*Iba1 staining in rTg^WT^ and rTg^ArcKO^ mice:* CA1 region of the dorsal hippocampus was imaged in 2 slices (2 images) for each animal. Imaging was performed through 40µm slices with 1µm step size. An overlay of Iba1 and DAPI was made, and the number of Iba1-positive cells were counted. Iba1 raw integrated density/area was calculated for each image. The values for each image were averaged to get a representative value for an animal.

### Statistical analysis

Unpaired t-test (with Welch correction), one-way ANOVA (with Sidak’s multiple comparisons test), and two-way ANOVA (with Tukey’s multiple comparisons test) were performed using GraphPad Prism (GraphPad Software, San Diego, CA). Column graph data points represent the mean ± SEM. All data sets were tested for normality and outliers. An alpha value of 0.05 was used to determine statistical significance for all tests (*p<0.05; **p<0.01; ***p<0.001; ****p<0.0001).

